# scCAT-seq:single-cell identification and quantification of mRNA isoforms by cost-effective short-read sequencing of cap and tail

**DOI:** 10.1101/2019.12.11.873505

**Authors:** Youjin Hu, Jiawei Zhong, Yuhua Xiao, Zheng Xing, Katherine Sheu, Shuxin Fan, Qin An, Yuanhui Qiu, Yingfeng Zheng, Xialin Liu, Guoping Fan, Yizhi Liu

**Affiliations:** State Key Laboratory of Ophthalmology, Zhongshan Ophthalmic Center, Sun Yat-Sen University, Guangzhou, China; Department of Human Genetics, David Geffen School of Medicine, UCLA, Los Angeles, CA, USA; Earth, Planetary and Space Sciences, UCLA, Los Angeles, CA, USA; Medical Scientist Training Program, David Geffen School of Medicine, UCLA, Los Angeles, CA, USA

## Abstract

The differences in transcription start sites (TSS) and transcription end sites (TES) among gene isoforms can affect the stability, localization, and translation efficiency of mRNA. Isoforms also allow a single gene different functions across various tissues and cells However, methods for efficient genome-wide identification and quantification of RNA isoforms in single cells are still lacking. Here, we introduce single cell Cap And Tail sequencing (scCAT-seq). In conjunction with a novel machine learning algorithm developed for TSS/TES characterization, scCAT-seq can demarcate transcript boundaries of RNA transcripts, providing an unprecedented way to identify and quantify single-cell full-length RNA isoforms based on short-read sequencing. Compared with existing long-read sequencing methods, scCAT-seq has higher efficiency with lower cost. Using scCAT-seq, we identified hundreds of previously uncharacterized full-length transcripts and thousands of alternative transcripts for known genes, quantitatively revealed cell-type specific isoforms with alternative TSSs/TESs in dorsal root ganglion (DRG) neurons, mature oocytes and ageing oocytes, and generated the first atlas of the non-human primate cornea. The approach described here can be widely adapted to other short-read or long-read methods to improve accuracy and efficiency in assessing RNA isoform dynamics among single cells.

## Background

The extent of cellular heterogeneity across different tissues and cell types has become increasingly apparent due to the development of genomics technology, especially single-cell omics sequencing (1-3). With the launch of initiatives such as the human single-cell atlas (4, 5), increased attention has been given to the regulatory mechanisms of cell-specific gene transcription, including both transcript abundance and alterative isoform usage, which can result in distinct protein sequences and structures (6, 7). RNA isoform variability includes intron inclusion, exon skipping, and alternative choice of transcription start sites (TSSs) (8) and transcription end sites (TESs) (9, 10). Alternative TSSs and TESs account for the majority of tissue-specific exon usage, are considered the principal drivers of transcript isoform diversity across tissues, and underlie the majority of isoform-mediated, cell-type specific proteomes (11). In addition, alternative TSS choices in the 5’-UTR, as well as alternative polyadenylation (APA) in the 3’-UTR regions play key roles in mRNA stability, translation, localization (9, 10, 12-14).

Previous studies have demonstrated the widespread heterogeneity of transcript isoforms with alternative 5’-TSS or 3’-APA across different cell types, resulting in the discovery of new transcripts with tissue- or cell-type specificity, and allowing updates to transcript annotations of reference genomes (13, 15). Despite considerable success in measurements made on bulk populations, current approaches for identifying RNA isoforms and the dynamics of TSS/TES choices in single cells are limited. Fundamentally, there is currently no method for accurate, efficient, and quantitative analysis of RNA isoforms of single cells genome-wide. Most single-cell transcriptome approaches are based on single-ended quantification of RNA molecules (5’ or 3’) which give partial information on one end but not the whole transcript (3, 16, 17), resulting in loss of important information about the other end, especially for transcripts regulated by UTR regions on both ends (13). Methods based on single-cell full-length cDNA amplification such as Smart-seq2 can detect the full-length cDNA, but its coverage at both ends is low, and it is not possible to accurately distinguish the start and end positions of different transcript isoforms of the same gene (18, 19). Recently, approaches based on long-read RNA sequencing technologies identified RNA isoforms of thousands of cells, but challenges still remain. For example, the sequencing depth needed to quantitatively assess the RNA isoform transcriptome makes long-read sequencing too expensive, and the conventional approach has been to first catalog isoforms using the long reads and then map short reads to the resulting transcriptome references for quantification. In addition, the requirement of several micrograms of cDNA input requires extensive PCR amplification from picograms of mRNA of a single cell, which unavoidably results in higher PCR bias towards specific isoforms.(13, 15, 20).

In order to address these problems, we developed a simple and efficient approach based on well-established short-read sequencing platforms to explicitly exploit transcription initiation and termination sites for the quantification of RNA isoforms in single cells. When deployed in conjunction with optimized machine learning models, scCAT-seq is more accurate and cost-effective, and has higher efficiency than existing methods, making it suitable for quantitative and qualitative analysis of isoform transcriptomes of single cells, and for analysis of RNA isoform dynamics in different biological contexts.

## Results

To develop scCAT-seq, we adopted a strategy to capture the boundaries of transcripts at both 5’ and 3’ ends (21). Full-length cDNAs were first tagged with specific sequences adjacent to both TSSs and TESs and further amplified, based on a modified Smart-seq2 protocol (19). Segments of transcript ends with sequence tags were then tagmented out by Tn5 transposases and captured by targeted PCR amplification. Illumina sequencing adaptors were further added for standard Illumina sequencing. As expected, the reads with tags are distributed at the terminal sides of transcripts (**Supplementary Fig. 1a, b**). The analysis pipeline precisely determined TSSs by the mapped position of reads with a head tag, along with the “GGG” signal added during reverse transcription. TESs were determined from paired reads (R1 containing a tail tag and R2 covering polyA sites) by mapping R2 sequences near polyA sites to the genome (**Fig. 1a**). Peaks were called using the CAGEr package (22). Internal TES peaks derived from the internal priming during reverse transcription of mRNA were excluded.

**Figure 1.**
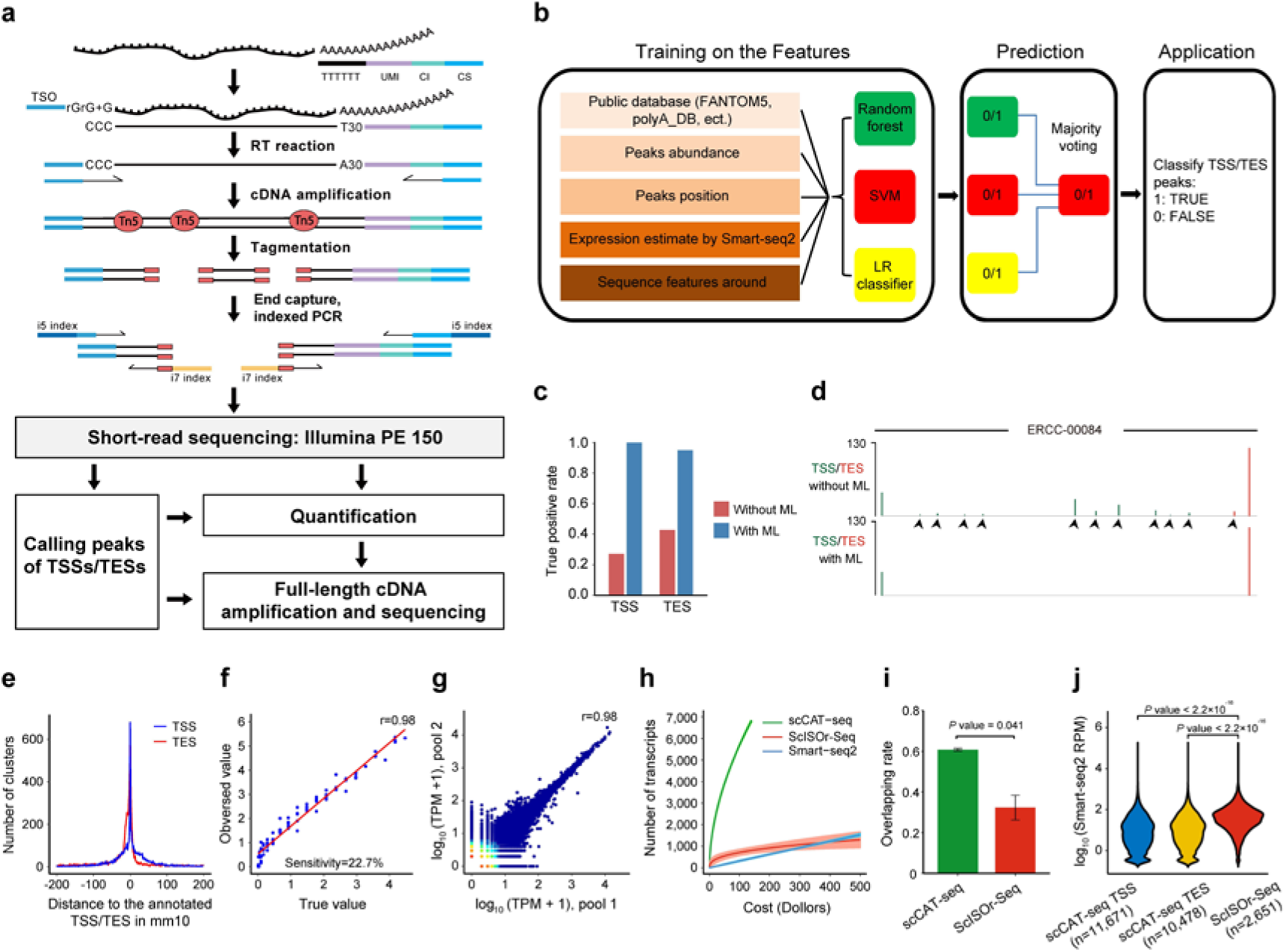
The scCAT-seq method and performance. **a**, Schematic of the scCAT-seq method. Template switching reverse transcription of full-length cDNA was performed with oligo-dT primer containing a unique molecular identifier (UMI), a cell identifier (CI), and a common sequence (CS). After PCR amplification, cDNA was tagmented with Tn5 transposases. Both 5’ and 3’ ends of the cDNA were captured and amplified by PCR using primers binding to CS and TSO sequences, during which Illumina sequencing indexes were tagged. In addition, Smart-seq2 libraries are generated from cDNA of the same cell. Sequencing data was processed and transcription start sites (TSSs) and transcription end sites (TESs) were identified by machine learning models, following by quantification of transcript isoforms. **b**, Schematic of the machine learning model. Features were collected and three machine learning models were implemented. Predictions from all models were integrated by majority voting. **c**, True positive for identification of TSSs and TESs with and without optimization by the majority voting strategy based on machine learning models. **d**, Genome browser shows the example of TSS/TES identification with or without machine learning (ML). The false positive peaks filtered out by ML were indicated by arrows. **e**, Distance of TSSs/TESs identified by scCAT-seq in the genome to those annotated in mm10. **f**, Scatter plot of observed transcript expression levels (y axis) and true abundance (x axis) of ERCC spike-ins through 5’-end quantification (n = 92). Each point represents a transcript. The Pearson’s correlation coefficient is shown in the upper right corner. The capture efficiency is estimated by the probability of an individual transcript could be detected at sequencing depth of one million. **g**, Scatter plots shows the Pearson’s correlation of transcriptional level of isoforms between replicated pools of 3 single cells. **h**, The number of transcripts with both ends captured using scCAT-seq, Smart-seq2, or ScISOr-Seq, versus cost. The shaded regions represent 95% confidence interval. **i**, Barplot shows the overlapping rate of genes detected among single cells, by scCAT-seq versus ScISOr-Seq (n = 3 single cells). Significance was computed using two sided t-test. Error bars represent standard error of the mean. **j**, Violin plot for expression level comparison between genes detected by scCAT-seq and ScISOr-Seq. Gene expression levels were quantified by Smart-seq RPM value. Significance was computed using two-sided Wilcoxon test.

To improve the accuracy in identification of real TSSs/TESs, we decided to employ machine learning models. Based on the read distribution of scCAT-seq and Smart-seq2 of the same single cell samples, we collected potential features that could affect the identification of a peak as a TSS or TES peak (**Table 1**), and implemented three widely used machine learning models: logistic regression classifier (LR), random forest (RF), and support vector machine (SVM). The random forest model indicated that “Slope_smart2_curve” and “Percentage” were the most important features, while the logistic regression classifier and SVM put the highest weights on “TPM_of_Dominant_Site” and “Trend_of_smart2_read_counts” (**Supplementary Fig. 1d, e**). To derive the best predictions, we chose an ensemble learning strategy of majority voting, integrating predictions from all models to systematically determine the real TSSs/TESs (**Fig. 1b**). Our strategy resulted in perfect performance on an independent test dataset of ERCC spike-in in single cells, with the true positive rate improved by 3.7- (27% versus 100%) and 2.2-fold (43% versus 95%) for TSS and TES, respectively (**Fig. 1c, d**), with sequencing depth of 4 million reads per sample (**Supplementary Table 1**). Similarly, ERCC data from other methods, such as C1 CAGE (17), C1 STRT (23), and a method developed by Arguel et al. (24), also showed high false positive rates for peaks identified as TSSs, (**Supplementary Fig. 1c**), but the accuracy was also improved to above 95% after using our machine learning model (**Supplementary Fig. 1f**), indicating that our model can be applied to other data sets that contain high false positive rates.

**Table 1.** Features used in the machine learning models.

|  | Features | Description of the features |
| --- | --- | --- |
| $x_1$ | TPM_of_peak | The total TPM value of the peak called by CAGEr. |
| $x_2$ | TPM_of_Dominant_Site | The highest TPM value of all sites within a peak. |
| $x_3$ | Gene_length | The length of the transcript annotated. |
| $x_4$ | Peak_width | The width of the peak called by CAGEr. |
| $x_5$ | Dominant_TPM_to_Smart2 | The ratio of Dominant_TPM to the RPM value of the corresponding gene revealed by Smart-seq2. |
| $x_6$ | Slope_smart2_curve | The slope of Smart-seq2 coverage curve around the peaks |
| $x_7$ | Trend_of_smart2_reads | Calculated by dividing the number of reads increased/decreased within 50bp distance by 50 |
| $x_8$ | Percentage | The percentage of read counts of a peak to the total counts of a transcript |

Using the sequencing data from mouse dorsal root ganglion (DRG) neurons for further benchmarking, we sequenced 18 DRG neurons with a mean of 2.4 million reads per cell (**Supplementary Table 2**). As genomic sequence features can specify the locations of TSSs and TESs, in addition to the eight features of read distribution, we added an additional 650 and 150 features of motifs related to TESs and TSSs. To train the TSS machine learning model, we used the data of neuron tissues from the FANTOM5 database (25), and to train the TES model, we used the mouse polyA sites peak from PolyA_DB (26). Using these databases, with 70% of the data for training and 30% for testing, we found the prediction accuracy for TSS and TES to be 94.3% and 94.2% respectively (**Supplementary Fig. 1g**). In total, after pooling all 18 cells together and applying the machine learning model, we identified 11991 and 15481 peaks as TSSs and TESs, which were significantly enriched at annotated TSS and TES regions, respectively (**Fig. 1e**). Over 93% of identified TSSs were located within 1 kb of annotated TSSs, and over 86% of identified TESs were within 1kb of annotated TESs (**Supplementary Fig. 1h, i**). In summary, our results indicate that scCAT-seq together with a machine learning model can identify TSSs and TESs of transcripts with high accuracy, allowing demarcation of transcription boundaries of full length isoforms.

Furthermore, we compared detected read counts with the known abundances of ERCC mRNA molecules to assess quantification performance. The measured abundances were highly concordant with the ground truth, with a Pearson’s correlation coefficient of 0.98 for both TSS and TES (**Fig. 1f, Supplementary Fig. 2a**). For the annotated genes of the mouse genome, an internal comparison between random pools of 3 single cells, each from the oocyte population, gave a correlation coefficient of 0.96 and 0.94 for the quantification of TSS and TES, respectively (**Fig. 1g, Supplementary Fig. 2b**). Thus, the quantification of TSS and TES is reliable and provides an accurate and reproducible measure of relative expression of transcript isoforms.

The sensitivity and efficiency were first estimated with ERCC spike-ins. The lowest detectable concentration was 4.4 molecules per million for both TSS and TES. In other words, at a detection threshold of TPM>1, at least 4.4 molecules are required to get one detected read at sequencing depth of one million. Therefore, the sensitivity of this method is estimated at roughly 22.7% (1/4.4) (**Fig. 1f**). This sensitivity is approximately the same as the 22%-26% sensitivity previously reported for detection of TSSs (24, 27, 28), but much higher than the 5.4% for TESs (29). In addition, the number of TSSs detected genome-wide by scCAT-seq is highly dependent on the number of reads mapped to the genome. Compared to existing methods which can detect only a single end of transcripts (either the 5’-TSS or the 3’-TES), scCAT-seq also has significantly better or comparable performance. When 1.28 million reads were mapped to the mouse genome, around 8000 transcripts were detected by scCAT-seq, comparable to the number for C1 CAGE (17) and the approach developed by Arguel et al. (24), but much higher than STRT-seq (21) and Smart-seq2 (19), which are the current single cell TSS profiling methods (**Supplementary Fig. 3a**). Similarly, for TES detection with 1.28 million uniquely mapped reads, scCAT-seq can determine TESs of more than 12000 transcripts, which is comparable to BAT-seq (29), and much higher than Smart-seq2 (**Supplementary Fig. 3b**). Further, we compared the performance of scCAT-seq to that of scISOr-seq (15, 20) which is the only method available for profiling the full-length transcript of single cells. We sequenced 6 single oocytes with the Pacbio sequel platform, with 54,000 circular consensus sequencing (CCS) reads per single cell (**Supplementary Table 3**), which is much higher than that of 270 reads per cell reported by Gupta et al. (15), and similar to that reported by Byrne et al. on the Nanopore platform (20). By normalizing the sequencing depth to the cost for both scCAT-seq and scISOr-seq, we found scCAT-seq had a much higher efficiency in capturing both ends of full length isoforms than scISOr-seq, 3122 versus 919 genes for scCAT-seq versus scISOr-seq at the equal cost for 4 million PE150 short-reads from Illumina (**Fig. 1h, Supplementary Fig. 3c**). Around 15% of the genes could be detected by both methods, with a higher overlapping ratio in highly expressed genes (**Supplementary Fig. 3d, e**). In addition, for the number of overlapping genes between single cells, scCAT-seq had a 2-fold higher overlapping ratio than scISOr-seq (60% versus 30%), highlighting the high consistency of scCAT-seq (**Fig. 1i, Supplementary Fig. 3f, g**). Comparison of the expression of the transcripts detected by scISOr-seq and scCAT-seq showed that scISOr-seq mainly detected the part of transcripts with the highest abundance (**Fig. 1j**), which only account for 1/4 of those detected by scCAT-seq. Furthermore, for the same coverage, our approach drastically reduces library preparation and sequencing cost. For instance, scCAT-seq only requires 1/73 of the cost required by scISOr-seq for 1000 transcripts covered (**Supplementary Fig. 2c**). These results indicate that scCAT-seq is a more cost-effective and reliable approach for quantitatively detecting both start sites and end sites of full-length transcripts at single cell level.

### Identification of novel transcripts with scCAT-seq

Leveraging the capacity to demarcate the boundaries a transcript, we set out to identify novel isoforms, both alternative TSSs/TESs of annotated genes and novel transcripts of unannotated genes (**Fig. 2a**). Data from mouse oocytes and DRG neurons was used for benchmarking. For annotated genes, we identified both alternative TSSs and TESs events, as evidenced by 3102 novel TSSs and 5746 novel TESs in oocytes (**Fig. 2b**), and 2031 novel TSSs and 4693 novel TESs in DRG neurons (**Fig. 2c**). In addition, 71 and 107 novel, unannotated transcripts were identified in DRG and oocytes respectively. Of note, many RNA isoforms identified by scCAT-seq, and validated by Smart2-seq and Sanger sequencing, were drop-out by scISOr-seq (**Fig. 2d, f, h**), indicating that scCAT-seq can identify novel transcripts with higher efficiency than scISOr-seq.

**Figure 2.**
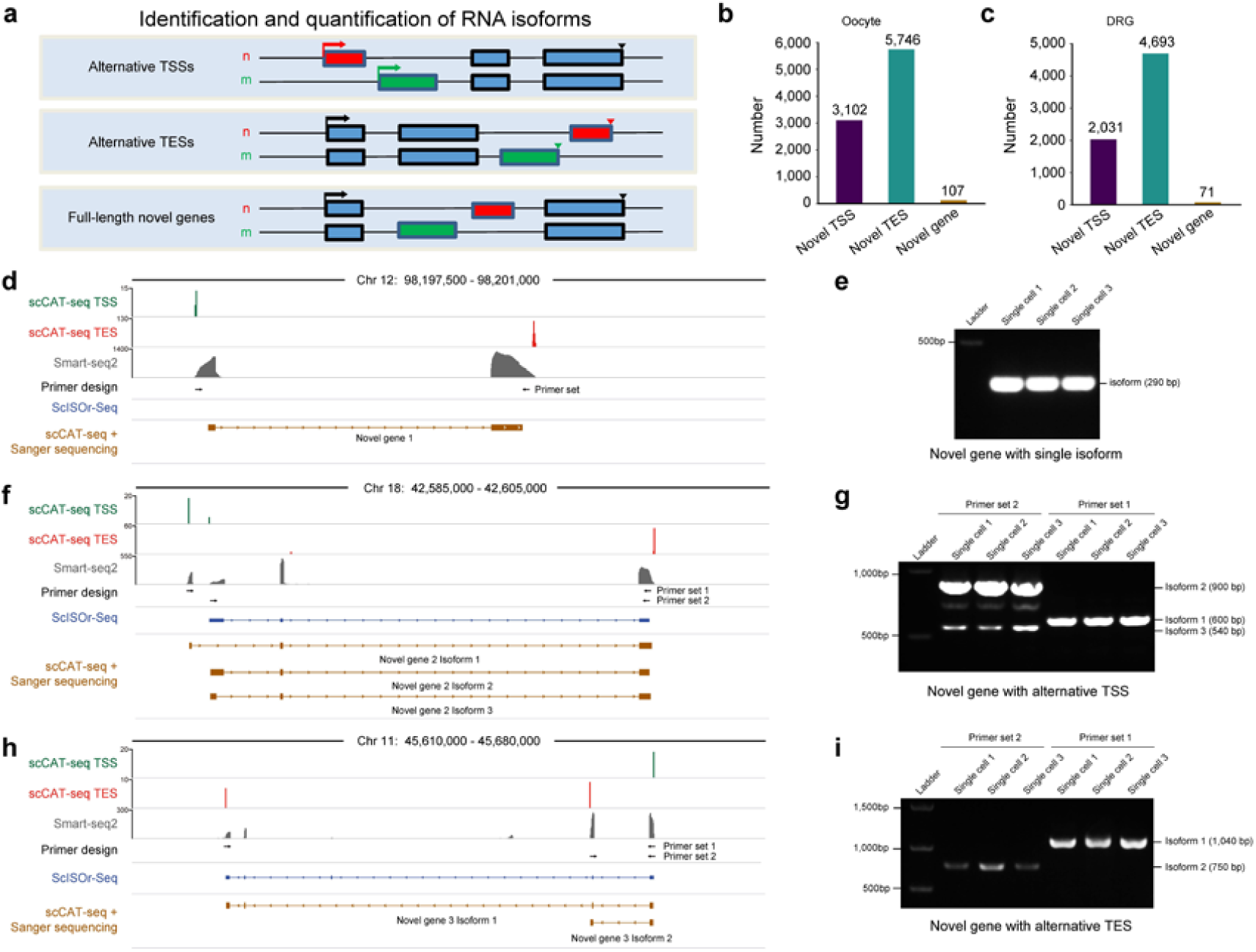
Characterization of novel transcripts and isoforms of single cells with scCAT-seq. **a**, Schematic of the functions of scCAT-seq. **b**, Barplot showing the number of novel isoforms of annotated genes and novel, unannotated transcripts in DRG neurons. The number of transcripts for each category is indicated above the box. **c**, Barplot showing the number of novel isoforms of annotated genes and novel, unannotated transcripts in oocytes. **d**, Genome browser track for an example of novel genes with single isoform. **e**, Gel image showing validation result of novel gene in **d. f**, Genome browser track for an example of novel genes with alternative TSSs on a different exon. **g**, Gel image showing validation result of novel gene in **f. h**, Genome browser track for an example of novel genes with alternative polyadenylation sites on a different exon. **i**, Gel image showing validation result of novel gene in **h**.

Further, to characterize the full-length information of novel RNA isoforms, such as alternatively spliced exons, full-length cDNAs were cloned with primers binding to the terminal ends identified by scCAT-seq (**Fig. 2a**). Full-length transcripts were sequenced by Sanger sequencing or scISOr-seq, and validated by Smart2-seq (**Fig. 2d-i**). For example, Figure 2f shows an example of novel gene with several isoforms, which were identified by Sanger sequencing of full-length cDNAs. Three isoforms differing in cDNA length have differential first exon choices (**Fig. 2f, g**), and alternative splicing events between isoform 2 and isoform 3 were revealed, which were also validated by Smart-seq2, including the exon not detected by scISOr-seq. In total, 96% (68/71) of novel transcripts detected by scCAT-seq were validated by Smart-seq2, while only 10% (7/71) of them were detected by scISOr-seq, indicating high drop-out rate of full-length transcripts in scISOr-seq. Our data suggest that when combined with targeted full-length sequencing, scCAT-seq can achieve higher coverage to reveal different isoforms of individual genes. In summary, scCAT-seq can accurately identify not only novel TSSs and TESs, but also completely unannotated full-length transcripts in single cells.

### scCAT-seq improves upon the performance of scISOr-seq for single cell RNA isoform quantification

Due to the higher efficiency and lower cost of scCAT-seq compared to scISOr-seq for identifying alternative isoforms, we hypothesized that scCAT-seq could also improve upon performance of scISOr-seq for accurately quantifying alternative isoforms (**Supplementary Fig. 4a**). It is currently too expensive to use scISOr-seq to obtain the sequencing depth required for accurate isoform quantification of multiple samples, especially at single cell level. Byrne et al. also tried to quantify isoforms with the number of CCS reads, but the number of genes covered was very limited. Concordantly, our data showed that the CCS readout for the majority of genes covered was less than 3 even though the sequencing depth was 0.5M for one single cell (**Supplementary Fig. 4b**). Although CCS read numbers are positively correlated with the number of reads of scCAT-seq, much higher variation was observed for the former with 10- to 1000-fold fewer read counts (**Supplementary Fig. 4c, d**). Intriguingly, when using the scCAT-seq to quantify the isoforms identified by scISOr-seq, the squared coefficient of variation (CV^2^) was reduced at least 10-fold, making isoform quantification much more accurate (**Supplementary Fig. 4d**). For example, two alternative isoforms of *Ermp1* were quantified with a CCS number below 5 in both DRG and oocytes, without sufficient power to differentiate the quantification of the two isoforms (**Supplementary Fig. 4e, f**). However, when quantified with scCAT-seq, with much lower variance, the longer isoform was found to be significantly higher expressed in oocytes than in DRGs. In summary, scCAT-seq can be used to quantify isoforms identified by scISOr-seq in single cells to improve accuracy with lower cost.

### Characterization and quantification of cell-type specific transcripts with scCAT-seq

To further assess differential gene expression between different cell types based on quantified abundances of TSS and TES tag counts, we performed scCAT-seq on three different cell types – mouse DRG, oocytes at Day 3, and oocytes at Day 4. Both TSS and TES transcriptome data clearly discriminated different cell types from each other (**Fig. 3a, Supplementary Fig. 5a**). In addition, because our method can identify both ends of transcripts, we set out to identify cell type specific transcript isoforms. Comparing DRG and oocyte cell-type specific isoforms, we identified 166 transcript isoforms encompassing 83 genes that only differed in TSS choices, and 222 isoforms encompassing 111 genes that only differed in TES choices (**Fig. 3b, Supplementary Fig. 5b, c**). For example, *Tsc22d1* and *Grpe1* had no difference in total gene expression between DRG and oocytes, but the two isoforms of each gene were expressed in a cell-type specific manner (**Fig. 3c, Supplementary Fig. 5d-f)**.

**Figure 3.**
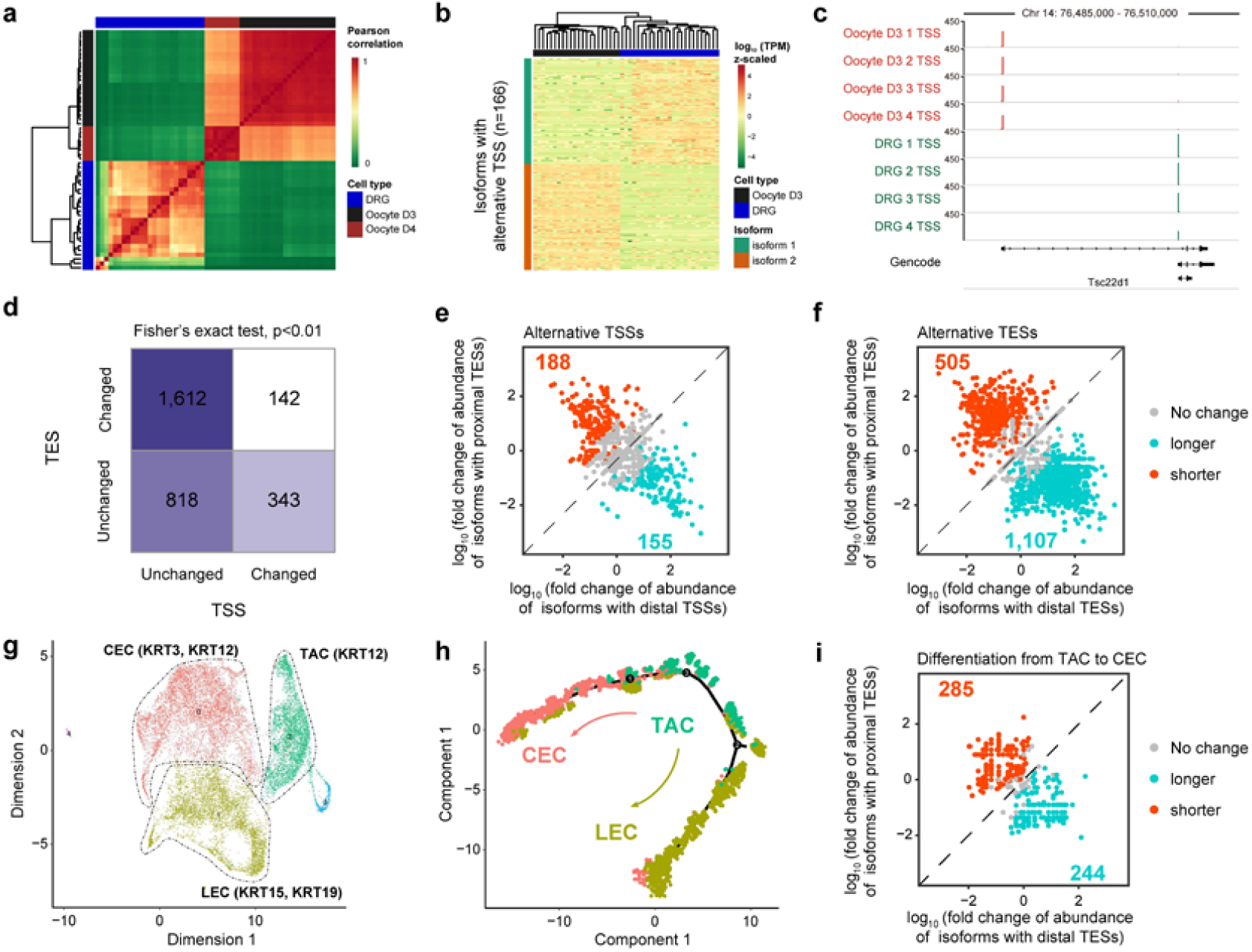
Quantification of cell specific isoforms discriminate cell types and illustrates the dynamics of isoform choices during oocyte ageing and corneal epithelial regeneration. **a**, Heatmap for Pearson’s correlation coefficient of transcriptomes of DRG neuron and oocytes, based on 5’-end quantification of RNA isoforms. **b**, Heatmap showing RNA isoforms of alternative TSS choices with cell type specificity. The major isoforms either in oocytes or in DRG neurons are shown (n = 166 isoforms). **c**, Genome browser tracks showing the alternative choices of TSS of *Tse22d1* between oocytes and DRG neurons. **d**, Heatmap showing the number of transcripts with or without TSSs/TESs changes during oocyte post-ovulatory ageing. **e**, Expression data with isoform specificity reveals isoform expression dynamics during oocyte post-ovulatory ageing (n = 1,161 genes). **f**, Expression data with isoform specificity reveals TES changes and isoform expression dynamics during oocyte post-ovulatory ageing (n = 1,754 genes). **g**, UMAP plot depicting cell clusters identified with scCAT-seq, including corneal epithelial cell (CEC), limbal corneal epithelial cell (LEC), and transient amplifying cell (TAC), and their specific marker genes (n = 7,848 cells). **h**, A pseudotime trajectory of single cells constructed using Monocle. Indicated in color are the three presumptive states corresponding to CEC, TAC, and LEC. **i**, Expression data with isoform specificity reveals TES changes and isoform expression dynamics during differentiation of CEC from TAC (n = 584 genes).

We also used scCAT-seq to assess RNA dynamics during ageing of post-ovulatory oocytes, and compared oocytes at day 3 post-ovulation (control) with oocytes at day 4 post-ovulation (post-ovulatory ageing oocytes). After assessing the 975 detectable TSSs and TESs across the control and ageing oocytes, we found that TESs are more prone to change positions, and the alternative choice of TESs is strongly associated with TSSs invariability (**two-sided Fisher’s exact test, P value = 3**.**0×10**^**-53**^), supporting the notion of interdependency between transcription initiation and polyadenylation (**Fig. 3d**). Further, a change in the choice of major isoform from day 3 to day 4 oocytes is observed in 343 genes with alternative TSSs and 1612 genes with alternative TESs, with a trend that shorter 5’ UTRs (**Fig. 3e**) or longer 3’ UTR are preferred (**Fig. 3f**). Thus, using scCAT-seq we can observe that the dynamics of major isoform choice during oocyte ageing is accomplished according to a general rule, which is through degradation of the major isoform on day 3, and activation of the minor isoform to switch to the alternative major isoform on day 4, as illustrated by *Ska3* (**Supplementary Fig. 6a**). In addition, the observations made by RT-qPCR validated our scCAT-seq data analysis (**Supplementary Fig. 6b**).

### Single cell atlas of non-human primate corneal epithelial based on RNA expression and APA analysis

We next employed scCAT-seq to profile a much larger number of single cells. Taking the non-human primate cornea as an example, we collected single cells and generated multiplexed cDNA using the 10x genomics platform. scCAT-seq libraries were subsequently generated and sequenced, and the 7848 single cells successfully captured were clustered into 5 major groups. Hundreds of marker genes for each cell type were identified (**Supplementary Fig. 7a**), with GO items relating to epithelial development enriched in the genes up-regulated and those relating to cell adhesion down-regulated (**Supplementary Fig. 7b, c**). Based on the RNA expression of the known marker genes, the following subtypes were identified: corneal epithelial cells (CEC) highly expressing KRT3 and KRT12, transient amplifying cells (TAC) highly expressing KRT12 but not KRT3, and limbal epithelial cells (LEC) highly expressing KRT19 (**Fig. 3g**). Pseudotime analysis on scCAT-seq data revealed the trajectory from TAC to LEC and CEC (**Fig. 3h**). We next identified the cell-type specific isoforms of the three major subtypes and assessed their dynamics. From TAC to LEC, we found 285 genes and 244 genes switched to proximal and distal APA sites, respectively (**Fig. 3i**). From TAC to CEC, we found 457 genes and 414 genes switched to proximal and distal APA sites, respectively (**Supplementary Fig. 8a**). For example, the longer isoform of UBE2B preferentially uses the distal TES in CEC, while the shorter isoform preferentially uses the proximal TES in TAC (**Supplementary Fig. 8b**). We also found that expression of genes with proximal APA sites was significantly higher in TAC than CEC/LEC, while there was no significant difference in expression between CEC and TAC for genes with distal APA sites in epithelial cells, suggesting a potential role of proximal APA choices in gene regulation during differentiation of epithelial cells from TACs (**Supplementary Fig. 8c-f**).

## Discussion

The approach we introduce here is highly accurate for transcript demarcation and isoform quantification in single cells. Through a machine learning algorithm that employs a majority voting strategy, the noisy false positive peaks were filtered out, enabling scCAT-seq to identify authentic terminal signatures with a true positive rate of 95%. Previously, machine learning has been successfully used to predict differential alternative splicing (30, 31), but none of them can be used to identify authentic demarcation of RNA isoforms to elucidate the transcriptomic complexity of single cells. The machine learning model developed here can also improve the accuracy of other methods to 95%, as evidenced by the ERCC data from C1 CAGE (17), C1 STRT (23), and Arguel et al., indicating that our model can be applied to other data sets that contain previously unrecognized high false positive signals. In addition to identification, the accuracy of our approach for quantification of the alternative isoforms is also very high, as the measured abundances are highly concordant with the ground truth, with a pearson’s correlation coefficient of 0.98. The high accuracy of both identification and quantification of isoforms provides an unprecedented opportunity for detection of previously unannotated genes and unidentified alternative TSSs and TESs, as well as for quantitation of cell-type specific RNA isoforms.

Another clear advantage of scCAT-seq is its efficiency. Based on short-read sequencing, scCAT-seq can identify TSSs and TESs simultaneously from sequencing data derived from a single library, enabling investigation of transcription initiation and polyadenylation in a large number of single cells. Compared with methods which capture only single ends of RNA transcripts, either the TSS or TES, scCAT-seq is demonstrably better for elucidating transcriptome complexity.

Compared with the recently developed long-read sequencing based method scISOr-seq, which can profile full-length transcripts for a group of single cells (15, 20), our approach requires 1/73 of the cost to detect the same number of transcripts, with higher efficiency. In addition, scISOr-seq requires at least 1ug of cDNA input, necessitating extensive amplification of cDNA with unavoidable PCR bias due the requirement for extra PCR cycles. This results in a decrease in the number of covered transcripts (a few hundred per single cell) and a lower transcript overlap ratio among single cells. In contrast, scCAT-seq only requires 0.1 ng of cDNA to achieve sufficient coverage of thousands of genes. Most importantly, it is still challenging to use scISOr-seq to quantify the isoforms differentially expressed between single cells, as accurate quantification requires deep sequencing that is currently too expensive for many labs. In contrast, our method can accurately quantify the transcripts (r=0.98) at an affordable cost for most labs. Due to the high accuracy and efficiency of scCAT-seq in identifying transcript ends, scCAT-seq also offers an efficient pipeline for full-length characterization of novel isoforms after targeted construction of full-length cDNA libraries, simply by PCR from the terminal sites identified by scCAT-seq in single cells.

In summary, the performance of scCAT-seq is a significant improvement upon that of scISOr-seq in terms of cost, efficiency, and accuracy of both identification and quantification of RNA isoforms.

Like all technologies, scCAT-seq has its limitations. First, the initial accuracy of TSS and TES identification is dependent on the effective cloning of full length cDNA. Although we adapted a widely used method Smart-seq2 to obtain cDNA, other protocols with better performance may be substituted in the future. Second, whereas the information of full-length isoforms of novel genes can be revealed by PCR using primers targeted to transcript ends identified by scCAT-seq, in this study we multiplexed only small number of example genes. However, profiling full-length transcripts with higher multiplexing can be done by complementing scISOr-seq downstream of scCAT-seq, in order to efficiently profile the targeted amplified full-length cDNA libraries. Including the scCAT-seq approach to initially identify isoforms of interest will help increase the efficiency of scISOr-seq with lower cost.

In conclusion, we believe that this robust and cost-effective approach is an ideal technology for comprehensive and systematic assessment of RNA isoform dynamics across heterogeneous single cells and biological conditions. Not only can it help define cell types with specific isoform expression patterns, but it can also establish a multi-faceted mammalian cell atlas in conjunction with other methodologies to identify tissue specific epigenetic elements, genotypes, and cis-elements. It can be widely implemented and may play important roles in projects such as the Human Single Cell Atlas.

## Methods

### Single cell isolation

The experiment was performed on 4-6 week old C57BL/6 mice of both genders. Mice were maintained under standard conditions (12 h light and dark cycles, with sufficient food and water). To obtain single DRG neurons, euthanasia was performed by CO_2_ and cervical dislocation, L4-L5 DRG from mice of both sides were dissected and dissociated into single cells. Single DRG neurons were manually picked by using a micro-capillary pipette. Single cells were incubated into a 0.2-ml thin-wall PCR tube containing 4 μl Smart-seq2 lysis buffer according to the published protocol(19, 32). To obtain postovulatory-aged oocytes, female mice were administered intraperitoneal injections of 10 IU pregnant mare serum gonadotropin and 10 IU human chorionic gonadotropin 48 hours later. Cumulus-oocyte-complexes (COCs) were collected 24 h after human chorionic gonadotropin injections from the oviductal ampullae. All cumulus cells were removed from the oocytes enzymatically by trypsin treatment (Sigma-Aldrich) for 2 min and oocytes were subsequently washed in DMEM medium containing 10% fetal bovine serum (FBS) (Sigma-Aldrich). Oocytes were picked into a 0.2-ml thin-wall PCR tube contains 4 μl Smart-seq2 lysis buffer as described before.

### scCAT-seq library construction

The full-length cDNA was generated through reverse transcription with transcriptase III and the RT primer (5’-AAGCAGTGGTATCAACGCAGAGTN4 [16bp of cell barcode] T30VN-3’), followed by PCR amplification according to Smart-seq2 protocol(19) with minor modification that Superscript II was replaced by superscript III to improve the yield of cDNA. ERCC RNA spike-in Mix which contains 92 transcripts (Thermo Fisher) was added and processed in parallel with poly-A RNA. After purification, 0.1 ng cDNA was used for Nextera tagmentation and fragments of both ends of the cDNA were selectively amplified by using the primers targeting TSO and Tn5 adaptors as shown in Fig. 1a. Library are purified using 1.8 × Agencourt AMPure XP beads (BECKMAN COULTER), and then loaded on an E-Gel 2% SizeSelect, and fragments of a length of 200-300bp bases were selected. Simultaneously, 0.1 ng of cDNA was used for standard Smart-seq2 libraries. Library was assessed by using Agilent Bioanalyzer 2100, and sequenced on Illumina Xten platform. The rest of the cDNA were used for PacBio ISO-seq analysis.

### Single cell ISO-seq

Single cell ISO-seq was performed on PacBio Sequel platform. Full-length cDNA of eight single cells were mixed together to reach the total amount of 2ug for each flowcell. PacBio library construction is done by using SMRTbell Template Prep Kit (PacBio cat#100-991-900), and sequenced using SMRTcells (PacBio cat#101-008-000), with eight single sample per SMRTcell.

### Single cell isolation of crab-eating monkey cornea epithelium and library construction

Whole eyes were dissected from a healthy crab-eating monkey. The lens, retina, iris, and trabecular network were removed and most of the conjunctiva was dissected and discarded. The corneal rims were subsequently treated with 1.5mL of 10mg/mL Dispase II in PBS at 37°C for 2 hours and 0.25% trypsin and 1 mM EDTA solution at 37°C for 15 minutes with gentle pipetting to yield single cells suspension. The disassociated corneal epithelial cells were captured on the 10x Genomics Chromium controller according to the Chromium Single Cell 3’ Reagent Kits V2 User Guide (10x Genomics PN-120237). Library was prepared using 0.3ng cDNA from 10x Genomics following the scCAT-seq protocol as described above.

### Data processing of next generation sequencing data

TSS and TES raw data were extracted and processed separately. For TSS data, reads with the sequencing tag 5’-GTGGTATCAACGCAGAGTACATGGG-3’ were selected, and TSO sequences 5’-GTGGTATCAACGCAGAGTACAT-3’ were trimmed away with the “GGG” tag retained. Then, these reads were aligned to mouse genome (mm10) with STAR (version 2.6.1a) with parameters (--outFilterMultimapNmax 1 --outFilterScoreMinOverLread 0.6 --outFilterMatchNminOverLread 0.6). Uniquely mapped reads were kept but discarded if the 5’ GGG was mapped. Reads that aligned to ribosomal RNA region were also discarded.

For the TES data, we first processed to remove 3’ adaptor sequences with cutadapt (version 1.18), and then extract pairing reads with R1 has 3’ Tag and R2 contains at least 10 polyA sequences at the 3’ side. Poly A sequences in the end of R2 were further trimmed with 5 A bases left at the 3’ side. By using STAR with parameters described above, reads were aligned to mouse genome (mm10). The reads with the terminal 5 A bases not mapped to the genome were retained for downstream analysis for polyadenylation sites. Reads mapped to multiple sites, with low quality alignment, and aligned to mitochondrial or ribosomal RNA region were discarded.

For Smart-seq2 data, raw reads past quality control were aligned by STAR using parameter as described above. Only reads that uniquely mapped to mm10 were retained and read count on each gene in each sample was computed using featureCounts (33). Differentially expressed gene analysis was performed using SCDE (version 2.10.1) (34).

For comparison, we downloaded BAT-seq data (accession number: GSE60768), C1 STRT (accession number: GSE60361) data and data generated by Arguel et al. (accession number: GSE79136) from the Gene Expression Omnibus database. C1 CAGE data were downloaded from DDBJ (Project ID: PRJDB5282). For the BAT-seq data, we picked 192 mouse ES cells. For the C1 STRT data, 80 mouse cerebellum cells from the single-cell dataset were randomly picked. Same strategies were used with small modification to process C1 STRT data and BAT-seq data. For all data, we converted bam files to bed files with BEDtools (version 2.27.1). For 5’ end data, we extract the 5’ end from bed files for further analysis. Likewise, we extract the 3’ end from bed files for 3’ end data.

### Data processing of scISOr-Seq data

Circular consensus reads (CCS) were obtained from the raw data of subreads Bam files by using PacBio Sequel SMRT-Link 7.0 Soft, with the default setting of parameters: minLength 10, maxLength 21000, minReadScore 0.75, minPasses 3. Then, reads were considered FLNC if they contained 5’ and 3’ primers in addition to a polyA tail. Primer and polyA tails were removed by cutadapt. Further, FLNC reads were aligned to reference genome mm10 using Minimap2(35) (version 2.17) with parameters (-t 30 -ax splice -uf --secondary=no -C5 -O6,24 -B4). CCS count on each gene in each sample was computed using featureCounts. The output Sam files were fed into Cupcake ToFU to collapse the mapped FLNC reads into unique transcripts. Scripts are available at: https://github.com/Magdoll/cDNA_Cupcake. Eventually, isoforms were identified and filtered using SQANTI2 against mm10 transcriptome annotation.

### Peak calling

To identify TSSs and TESs, we used CAGEr (version 1.24.0) package in R. Peaks were called using distclu (threshold = 5, nrPassThreshold = 1, thresholdIsTpm = TRUE, removeSingletons = FALSE, keepSingletonsAbove = 10, maxDist = 20). The position of dominant TSS/TES in each peak was set to represent the position of peak. TSS and TES annotation reference was based on gencode release_M18, and peaks mapped between 2kb upstream the annotated TSSs and 2k downsteam the annotated TESs were considered to belonging to the said gene. We then extracted 5’-end and 3’-end of all annotated transcripts and converted to bed files with a custom R script, and distance between the called peaks and the nearest annotated TSS/TES was calculated by a custom script. We adopted the following priority in calculating the distribution of TSS peaks mapped to genome features: TSS±1000 > 5’ UTR > first exon > first intron > other exon > other intron > 3’ UTR > intergenic. Similarly, The priority in calculating distribution of TES peaks mapped to genome features is TES±1000 > 3’ UTR > last exon > last intron > other exon > other intron > 5’ UTR > intergenic.

### Machine learning analysis

To predict a peak is real or false TSS/TES, we employed three widely used models, including logistic regression classifier, random forest and support vector machine.

Firstly, we use eight features of read distribution to train the three machine learning models and they are summarized in the table 1. To perform the analysis, we used two independent data sets derived from ERCC spike-ins which can serve as a standard for true TESs/TSSs determination, one is for training data, and the other is for test data. The features were normalized, “TPM_of_peak”, “TPM_of_Dominant_Site” are firstly being taken a log and secondly normalized to be in the range of [0,1]. Training data was generated by using scCAT-seq for ERCC spike-ins, and the test data was derived from the ERCC spike-ins mixed in the single cells. The True positive and False positive of TSSs and TESs predicted was calculated. Secondly, for the genomic data, in addition to the eight features of read distribution, we added an additional 650 and 150 features of motifs related to TESs and TSSs, and used FANTOM5 database (25) and PolyA_DB (26) to train the model for TSS and TES prediction respectively. TSSs/TESs were predicted from the peaks of single cells based on scCAT-seq. We utilize the popular open source python machine learning library scikit-learn to train these models.

With a logistic regression model, the probability () of a peak with the given values of the features (x_1,_ x_2,_ x_3,_ x_4,_ x_5,_ x_6,_ x_7,_ x_8_) was determined as:

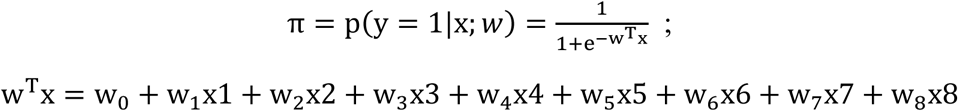

Where the (x_1,_ x_2,_ x_3,_ x_4,_ x_5,_ x_6,_ x_7,_ x_8_) are the observed value of the features shown in **Table 1**, and w_0,_ w_1,_ w_2,_ w_3,_ w_4,_ w_5,_ w_6,_ w_7,_ w_8_ are the coefficients of the corresponding features of the training model. The decision was made based on the following function:

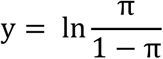

We also applied l2 regularization and the coefficient is determined using cross validation on the training set.

Random forest model(36) consists of a large number of individual decision trees that operate as an ensemble. Every tree in the random forest makes its own class prediction and the class with the most votes becomes the random forest model’s prediction. In random forest, each decision tree is independently trained using partial features and bootstrap sampled training data. To generate a tree, it has to go over each feature, and find the best one has the maximum gini index reduction(37) after splitting. The gini index for each node is defined as:

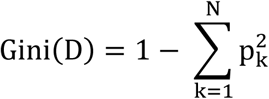

Where D is a node in the tree, N is the number of different classes, and p_k_ is the percentage of data in this node that is labeled class k. Conceptually, gini index reflect how different they are if we randomly choose two samples from the node. The smaller the gini index, the more pure the node is. After the split, if the child node still contains more than one class, it will go through the search process again to split it. This process generally ends when all the leaf nodes contain only one class samples. An example of a decision tree learned is shown as below:

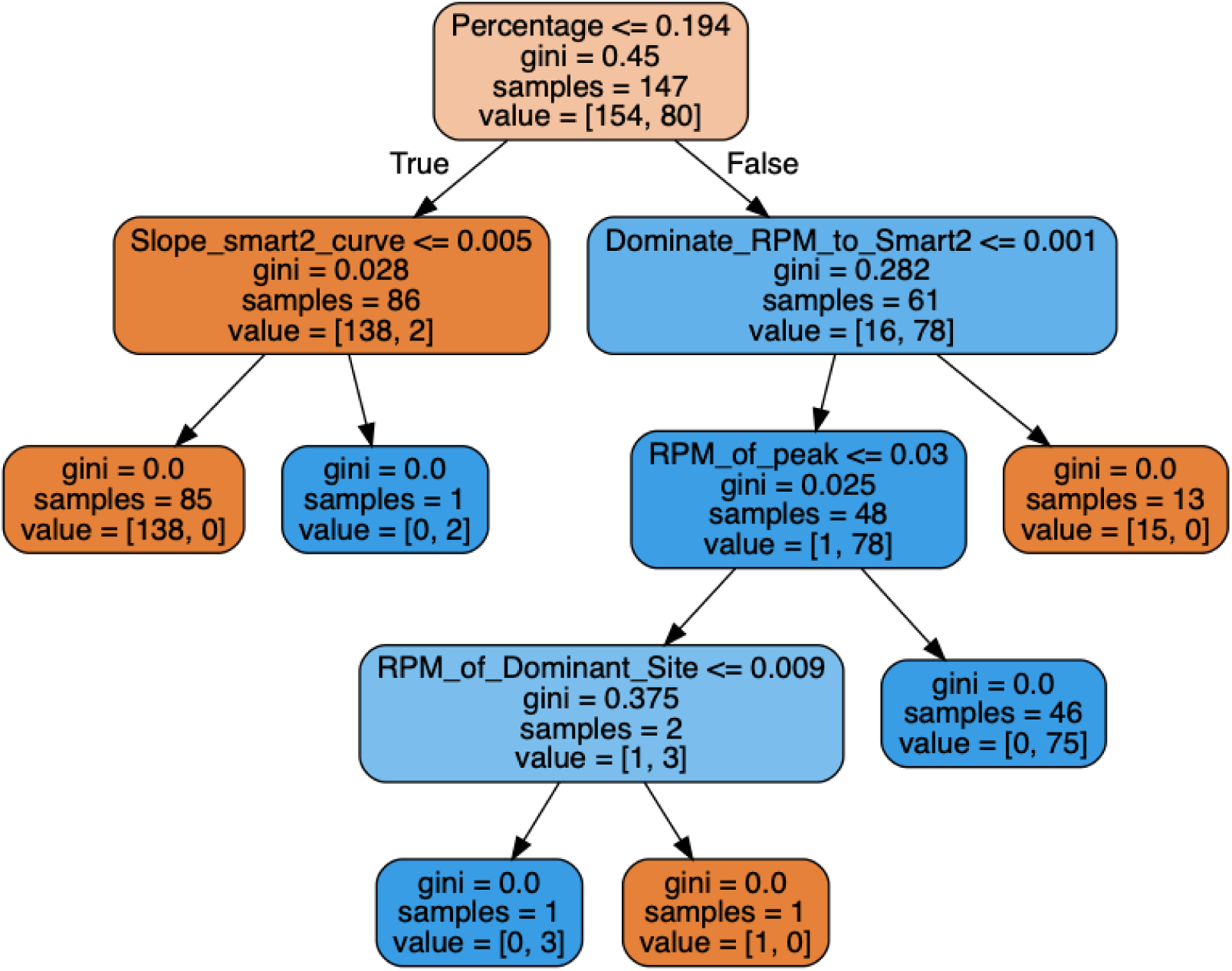

In every node of the tree in this plot, it first shows the selected feature and splitting criteria to maximize gini index reduction. Then it shows the gini index for this node. The “samples” represents number of distinct samples this node has. And the value has two numbers, corresponding to the number of negative and positive training data (some training data can have multiple copies since they are bootstrapped from the original dataset). After we have learned a number of decision trees, we’ll do a majority vote using all the trees’ predictions. In statistical theory, this step helps reduce model variance.

The SVM(38) is another widely used supervised machine learning models for two class classification (can be extended for multi-class classification and regression as well.) The SVM algorithm tries to find a hyper plane in a mapped high dimensional space (with kernel trick) that separates the two classes that achieves largest margin. From any textbook, the SVM with soft margin and regularization is formularized as:

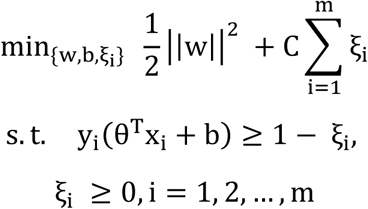

Where *ξ*_*i*_ is used to allow soft margin, and m is the number of training data you have. The C controls the relative regularization and is determined using cross validation method. And *w* is the vector of weights, and x is the feature vector.

Lastly, we try to further improve model performance by ensemble all three models. Dietterich (39) indicated statistical, computational and representational benefits of combining models. This theory is also validated here as the ensembled model achieves better performance than any one of the three models alone, despite the fact that the three models already achieve good performance on their own.

### Quantification of cell-type specific isoforms

Expression values for each peak (TSS/TES) were quantified as tags per million (TPM) generated by CAGEr. To identify cell-type specific isoforms, the major TSS/TES positions of genes co-expressed between the two types of cells are compared by intersect the bed files of each with BEDtools (40). Genes with either alternative TSS or alternative TES between the two were selected. Then, the differential expression analysis on the TPM value of the major isoform of each cell type between the two was performed with DESeq2. **Further**, we performed qRT-PCR with Unique Aptamer™ qPCR SYBR^®^ Green Master Mix (Novogene) on the RocheLightCycler480 (Roche) using the same samples used for next-generation sequencing to validate the alternative TES in different cell types. All assays were run in triplicate for six individual samples. The qRT-PCR conditions used were as follows: 5 min at 95°C, 45 cycles of 10 sec at 95°C and 30 sec at 60°C. The qRT-PCR primers sequences used were listed in Supplementary Table 4. Gene body primers were used to quantify total gene abundance. 3’ UTR primers were used to quantify long 3’ UTR isoform. Data were analyzed using the 2^-ΔΔCt^ method.

### Sequencing full-length cDNA of target genes

Primers were designed according to the coordinates of TSS/TES identified by scCAT-seq. Full-length cDNA of all isoforms of a target gene was amplified by PCR from the cDNA pool of single cells generated with Smart2-seq. Briefly, 1 ng full length cDNA was used to perform 35-cycle PCR with Premix Taq™ (TaKaRa). PCR products were purified with QIAquick Gel Extraction Kit (Qiagen) and Sanger-sequenced with corresponding primers. All assays were performed for three individual single cell samples. PCR primers used for novel genes are listed in Supplementary Table 5.

### Data processing of corneal single-cell data

Each 10x droplet sequencing data was processed using the Cell Ranger (version 2.1.1) pipeline from 10x Genomics. In brief, reads was demultiplexed and aligned to the *Macaca fascicularis* genome. UMI counts was quantified to generate a gene-barcode matrix. Cells were filtered to remove those containing less than 500 genes. Genes that were detected in less than 3 cells were also removed. Further analyses of these cells were performed using the Seurat (version 3.0.2) R packages, as described in the tutorials (“https://satijalab.org/seurat/”)(41). Briefly, cells were normalized using LogNormalize and multiplied by a scale factor of 10000. HVGs (high variable genes) were identified and used for further analysis. Shared cell states were identified using integration procedure in Seurat.

Dimensionality reduction was performed using principal component analysis (PCA). Statistically significant PCs were identified using the Jackstraw function. The score of cells in those significant PCs were used to build a k-nearest neighbor (KNN) graph. Louvain algorithm was used for identifying cell clusters in KNN graph (parameter resolution=0.06). Uniform manifold approximation and projection (UMAP) dimensionality reduction was used to project these populations in two dimensions. Pseudotime analyses of CEC was performed using Monocle2 (42) (version 2.12.0) R package. Differentially expressed genes among LEC, CEC and TAC were identified using differentialGeneTest function and used as input for temporal ordering of those cells along the differentiation trajectory.

## Code availability

Custom computer code used in this study is freely available at https://github.com/huyoujinlab/scCAT-seq.

## Availability of data and material

All the related data can be downloaded from GEO with the accession number **GSE134311**.

## Acknowledgement

The work is supported by National Key R&D Program of China (2018YFA0108300, 2017YFC1001300); National Natural Science Foundation of China (31700900; 81530028; 81721003); Clinical Innovation Research Program of Guangzhou Regenerative Medicine and Health Guangdong Laboratory (2018GZR0201001); Local Innovative and Research Teams Project of Guangdong Pearl River Talents Program; the State Key Laboratory of Ophthalmology, Zhongshan Ophthalmic Center, Sun Yat-sen University.

## Figure

**Supplementary figure 1.**
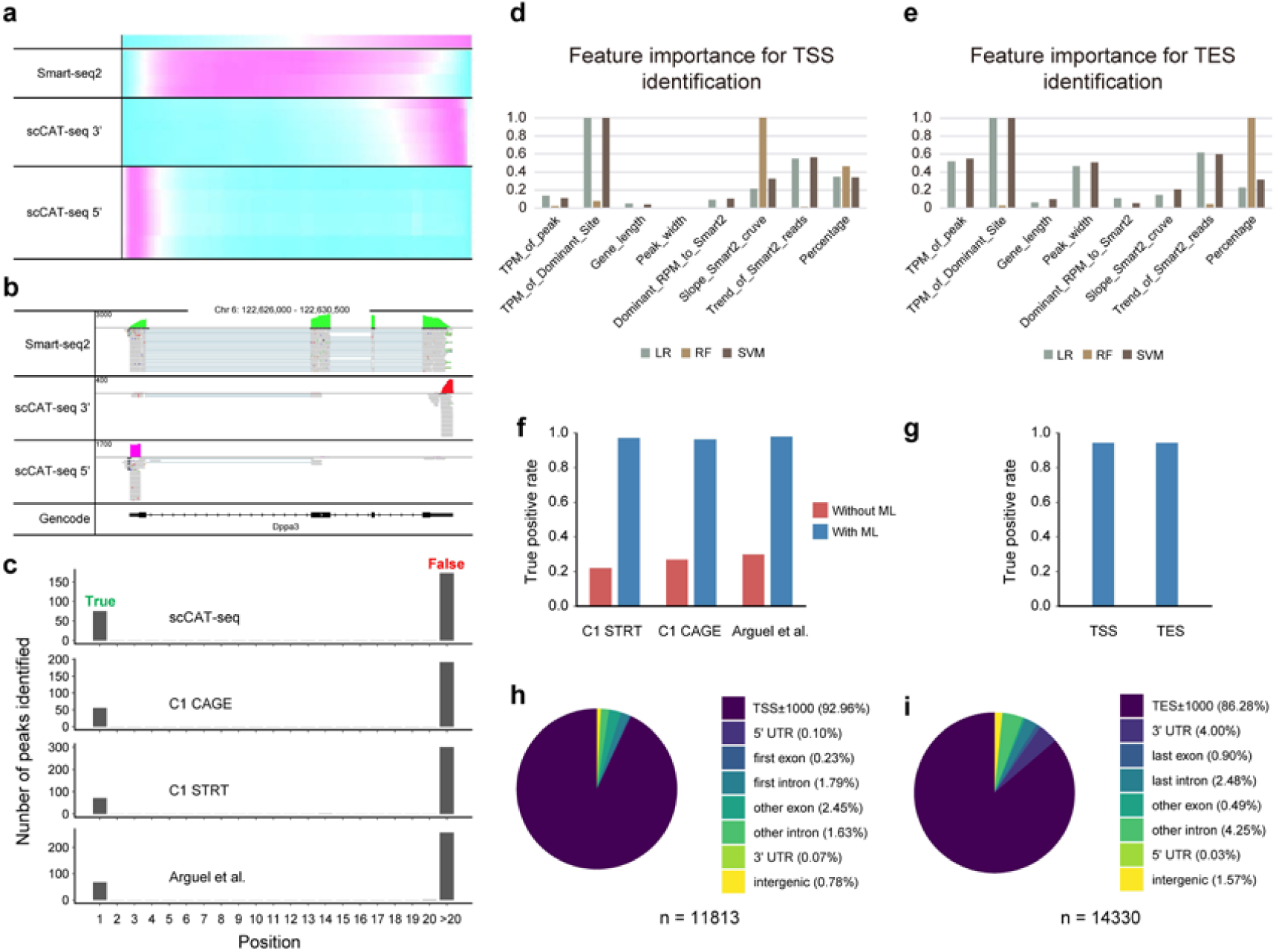
Machine learning improves accuracy of scCAT-seq demarcated isoform boundaries in single cells. **a**, Distribution of sequencing reads along the genes from head to tail from Smart-seq2 and scCAT-seq. **b**, *Dppa3* as an example gene, showing the distribution of sequencing reads of Smart-seq2 and scCAT-seq. **c**, TSS peaks identified in the data of scCAT-seq, as well as public data sets of ERCC for C1 CAGE, C1 STRT and Arguel et al. True positive peaks located around the annotated TSSs and false positive TSS peaks located elsewhere are indicated respectively. **d**, Relative feature importance of the eight features for TSS identification in random forest model (RF), support vector machine (SVM), and logistic regression classifier (LR). **e**, Relative feature importance of the eight features for TES identification in the three machine learning models. The value of importance for SVM and LR are first transformed to absolute value and normalized to the highest value of the eight features. **f**, True positive for identification of TSSs for the public data sets with and without optimization by the majority voting strategy based on machine learning models. **g**, True positive for identification of TSSs and TESs for the pooled single DRG neurons data sets with optimization by the majority voting strategy based on machine learning models. **h**, Pie chart with the genomic distribution of the identified TSSs. The total number of TSS peaks identified after optimization by the machine learning models is indicated under the pie chart. **i**, Pie chart with the genomic distribution of the identified TESs.

**Supplementary figure 2.**
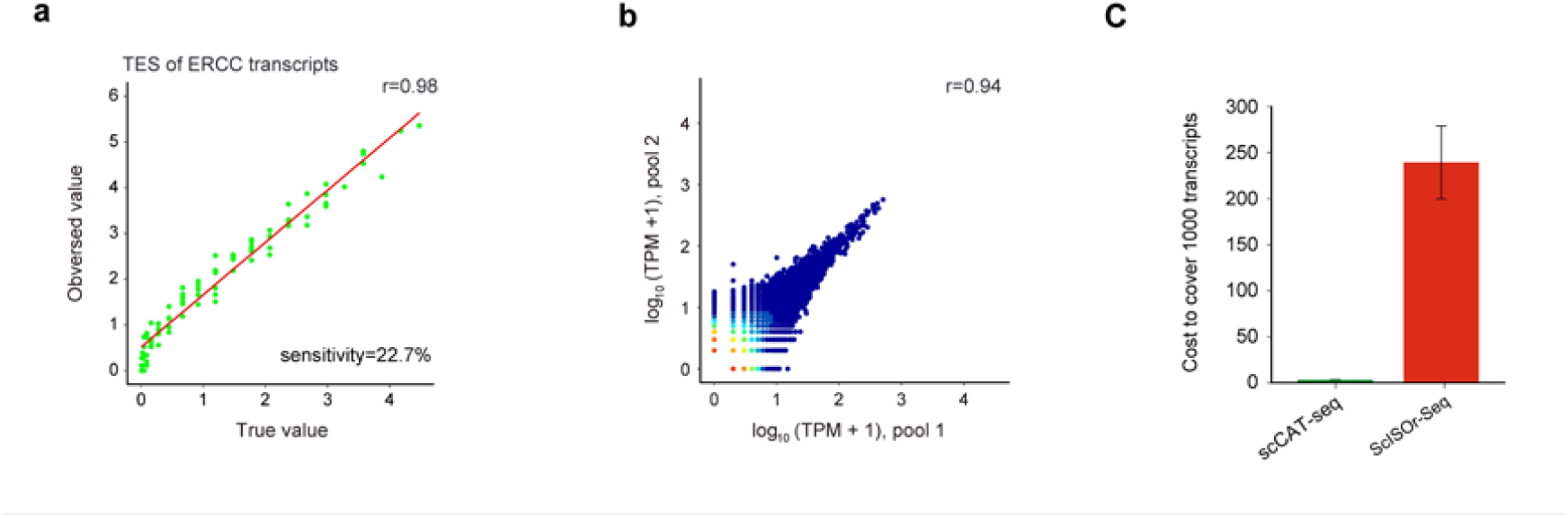
Accuracy and consistency of scCAT-seq performance for isoform quantification. **a**, Scatter plot of observed transcript expression levels (y axis) and true abundance (x axis) of ERCC spike-ins through 3’-end quantification. **b**, Scatter plots showing the correlation of transcriptional level of isoforms between replicated samples (3 cells pooled) based on 3’-end quantification. **c**, Comparison of the cost for the same number of transcripts (1,000) between PacBio ScISOr-Seq, scCAT-seq. The price is estimated based on the market price in China. Error bars represent 95% confidence interval.

**Supplementary figure 3.**
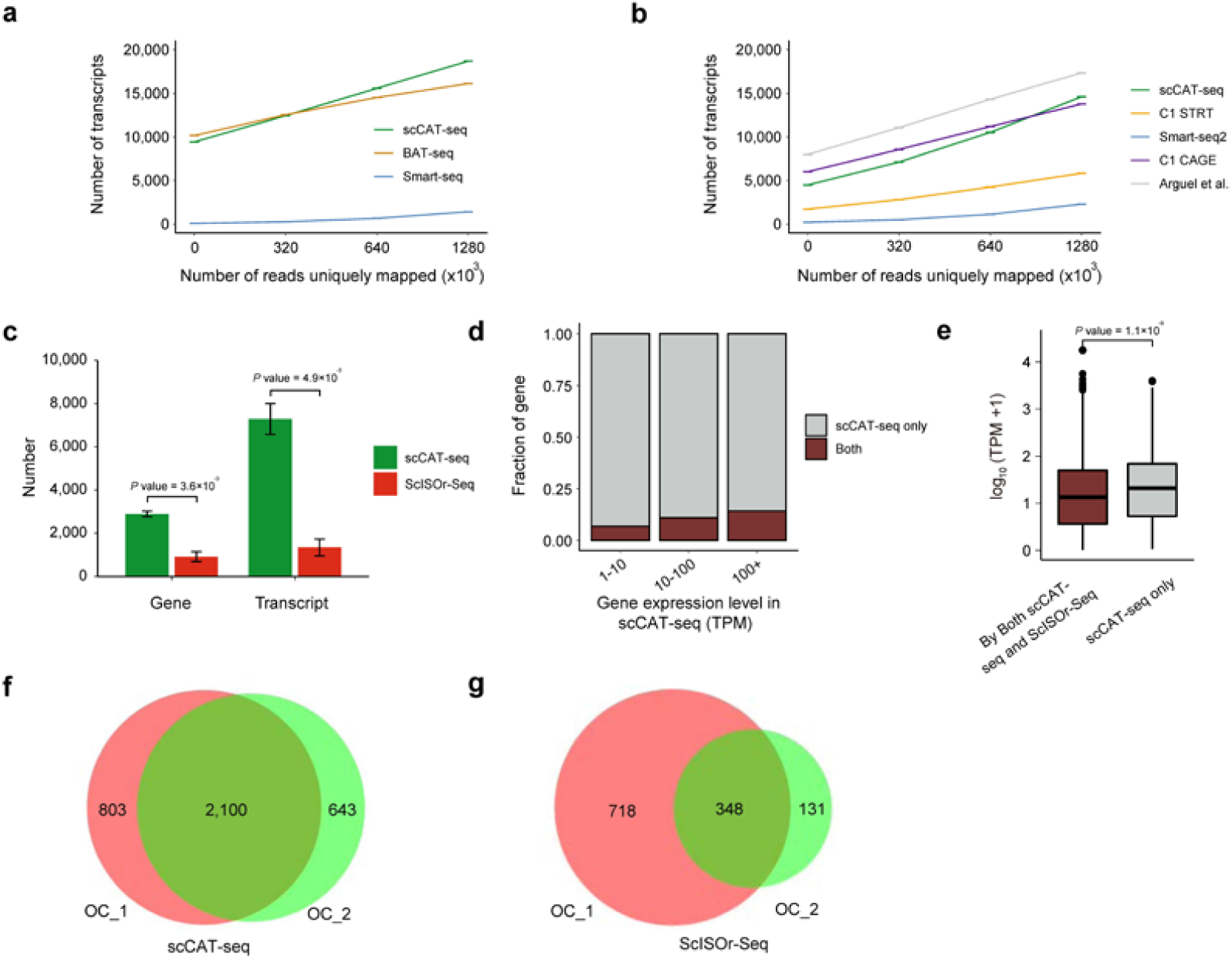
The number of genes covered by scCAT-seq. **a**, The number of transcripts with 3’ tail detected by scCAT-seq, BAT-seq, and Smart-seq2 at variable sequencing depth. Error bars represents standard error of the mean. **b**, The number of transcripts with 5’ head detected by scCAT-seq, C1 STRT, Smart-seq2, Arguel et al., and C1 CAGE at variable sequencing depth. Error bars represents standard error of the mean. **c**, Number of genes and transcripts covered by scCAT-seq and ScISOr-Seq respectively (n = 3). The number of reads for scCAT-seq was 4 million per single cell and the CCS number for ScISOr-Seq is 50,000 per cell. Significance was computed using two sided t-test. Error bars represents standard error of the mean. **d**, Stacked barplots showing the number of genes with different expression levels detected in oocytes by scCAT-seq and ScISOr-Seq. **e**, Boxplot for expression level comparison between genes detected by scCAT-seq only and by both scCAT-seq and ScISOr-Seq (n = 9,626). Significance was computed using two-sided Wilcoxon test. **f**, Venn diagram for genes detected concordantly among single cells by scCAT-seq. **g**, Venn diagram for genes detected concordantly among single cells by ScISOr-Seq.

**Supplementary figure 4.**
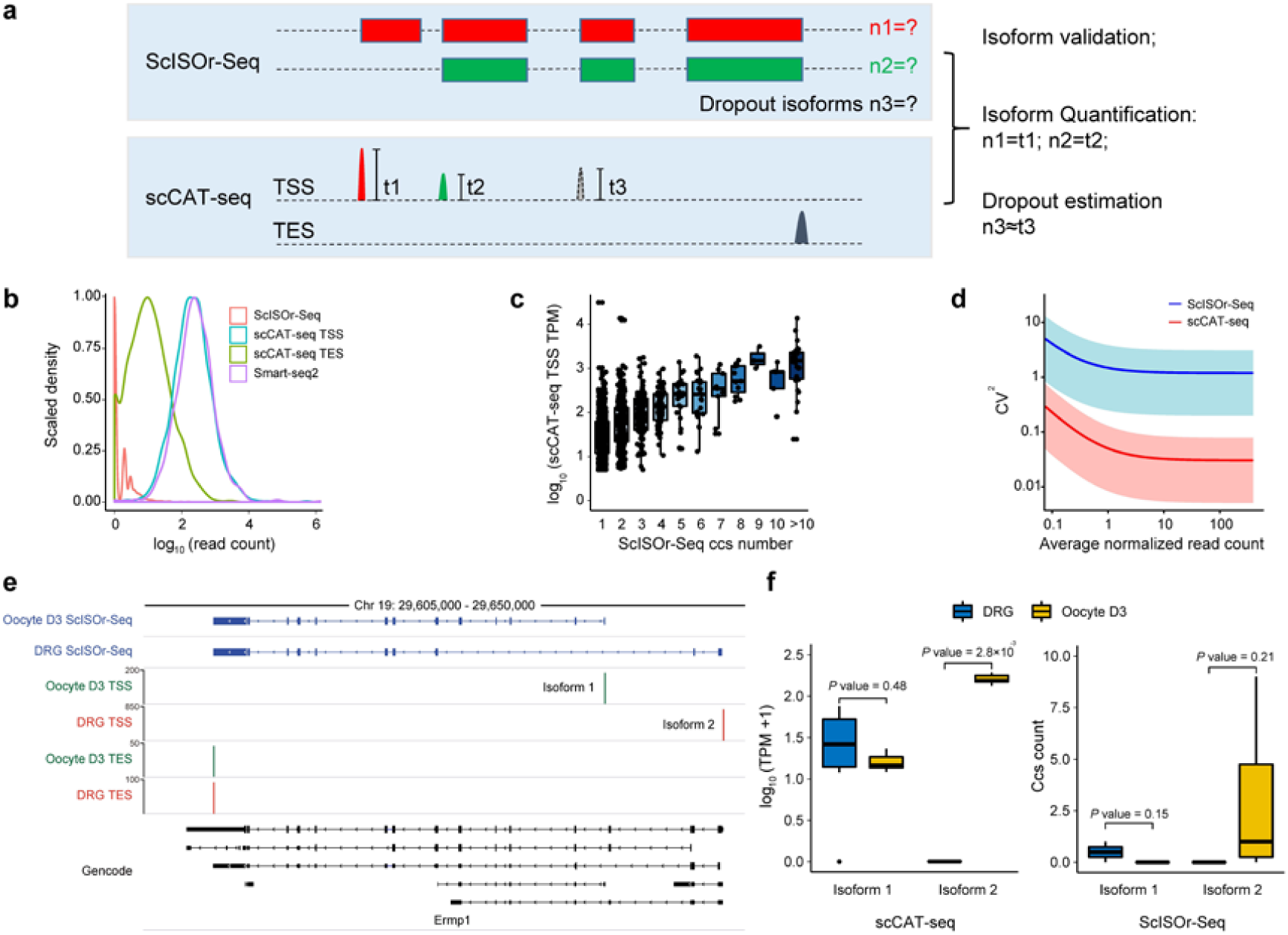
scCAT-seq improves upon the performance of scISOr-seq for single cell RNA isoform quantification. **a**, Schematic showing performance improvement of ScISOr-Seq via scCAT-seq quantification. **b**, Density plot showing the comparison of genes read count values among ScISOr-Seq, scCAT-seq and Smart-seq2. **c**, Boxplot for the expression level comparison among genes with different CCS numbers detected by ScISOr-Seq. **d**, Squared coefficients of variation of scCAT-seq and ScISOr-Seq, versus the means of normalized read counts. The shaded regions represent 95% confidence interval. **e**, Genome browser track showing an example of *Ermp1*. which has two isoforms detected by both scCAT-seq and ScISOr-Seq. **f**, Boxplot for the example gene *Ermp1*, which has two isoforms differentially expressed in oocytes or in DRG neurons, while the expression value assessed by ScISOr-Seq is not differential between the two cell types. Significance was computed using two-sided Wilcoxon test.

**Supplementary figure 5.**
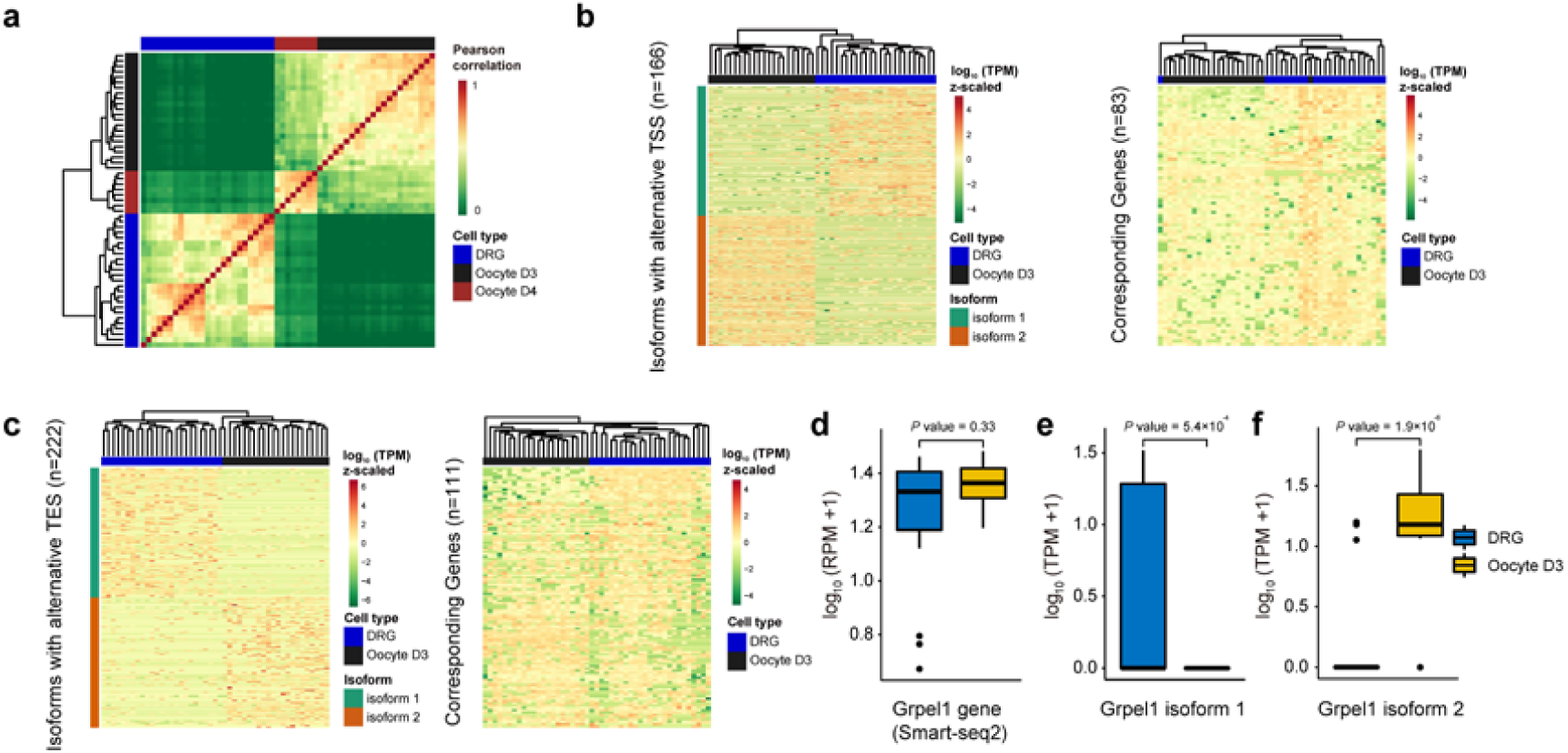
Identification and quantification of cell-type specific transcript isoforms. **a**, Heatmap for Pearson’s correlation coefficient of transcriptomes of DRG neuron and oocytes, based on 3’-end quantification of RNA isoforms. **b**, Heatmap showing RNA isoforms of alternative TSS choices with cell type specificity (left panel), and the expression of corresponding genes assessed by Smart-seq2 (right panel). **c**, Heatmap showing RNA isoforms of alternative TES choices with cell type specificity (left panel), and the expression of corresponding genes assessed by Smart-seq2 (right panel). **d-f**, Boxplot for the example gene *Grpe1*, which has two isoforms differentially expressed in oocytes or in DRG neurons (**e, f**), while the overall gene expression assessed by Smart-seq2 is not differential between the two cell types (**d**). For **d-f**, significance was computed using two-sided Wilcoxon test (n = 35).

**Supplementary figure 6.**
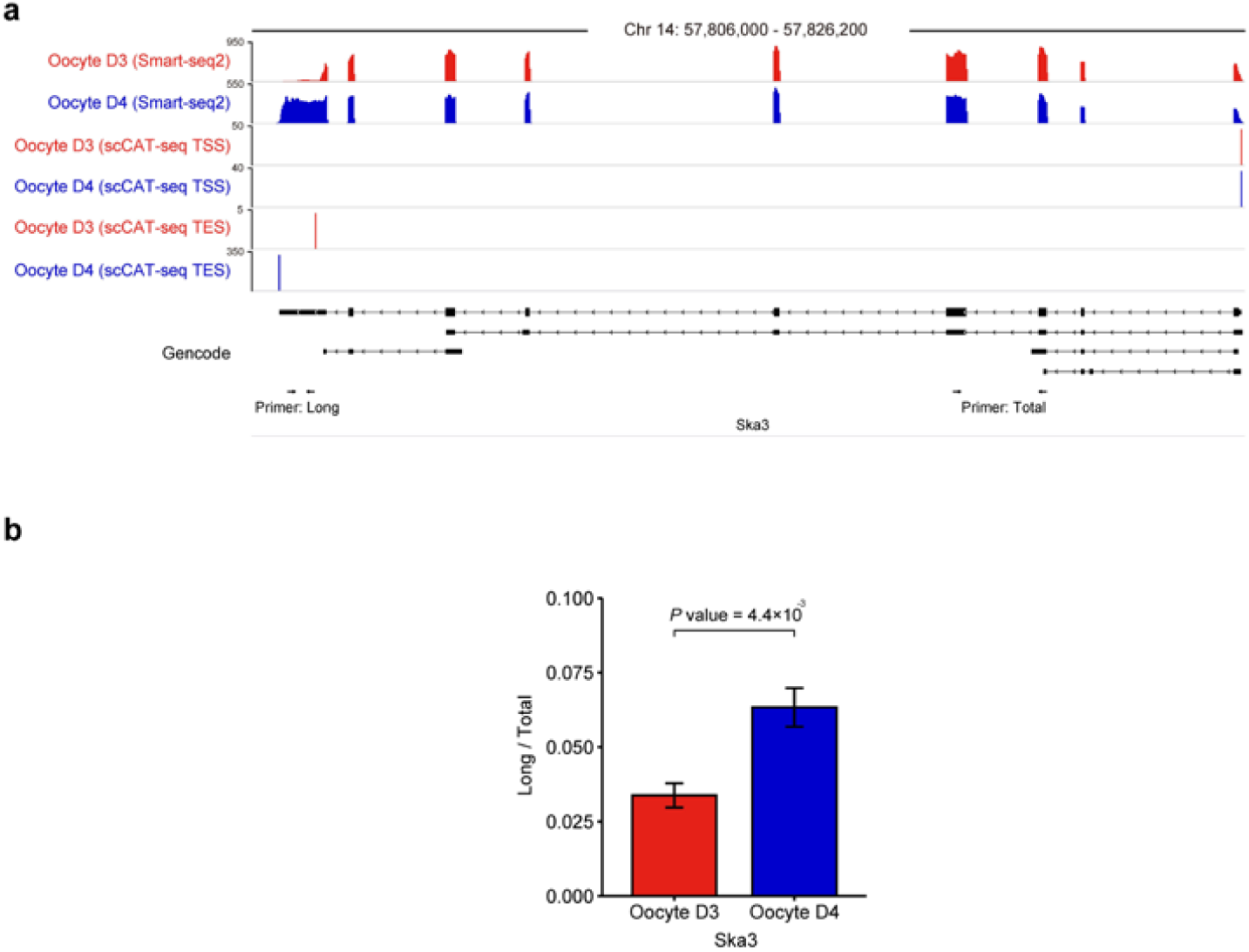
Examples of full-length isoforms with alternative TES during oocyte post-ovulatory ageing. **a**, Genome browser track showing the TSS dynamic choices during oocytes post-ovulatory ageing. **b**, Fold change in expression of the *Ska3* long 3’ UTR isoform (long) relative to total *Ska3* expression (total) between oocyte D3 and oocyte D4 single cells, measured by RT-qPCR. Error bars represent standard error of the mean (n = 6 biological replicates). Significance was computed using two-sided t-test.

**Supplementary figure 7.**
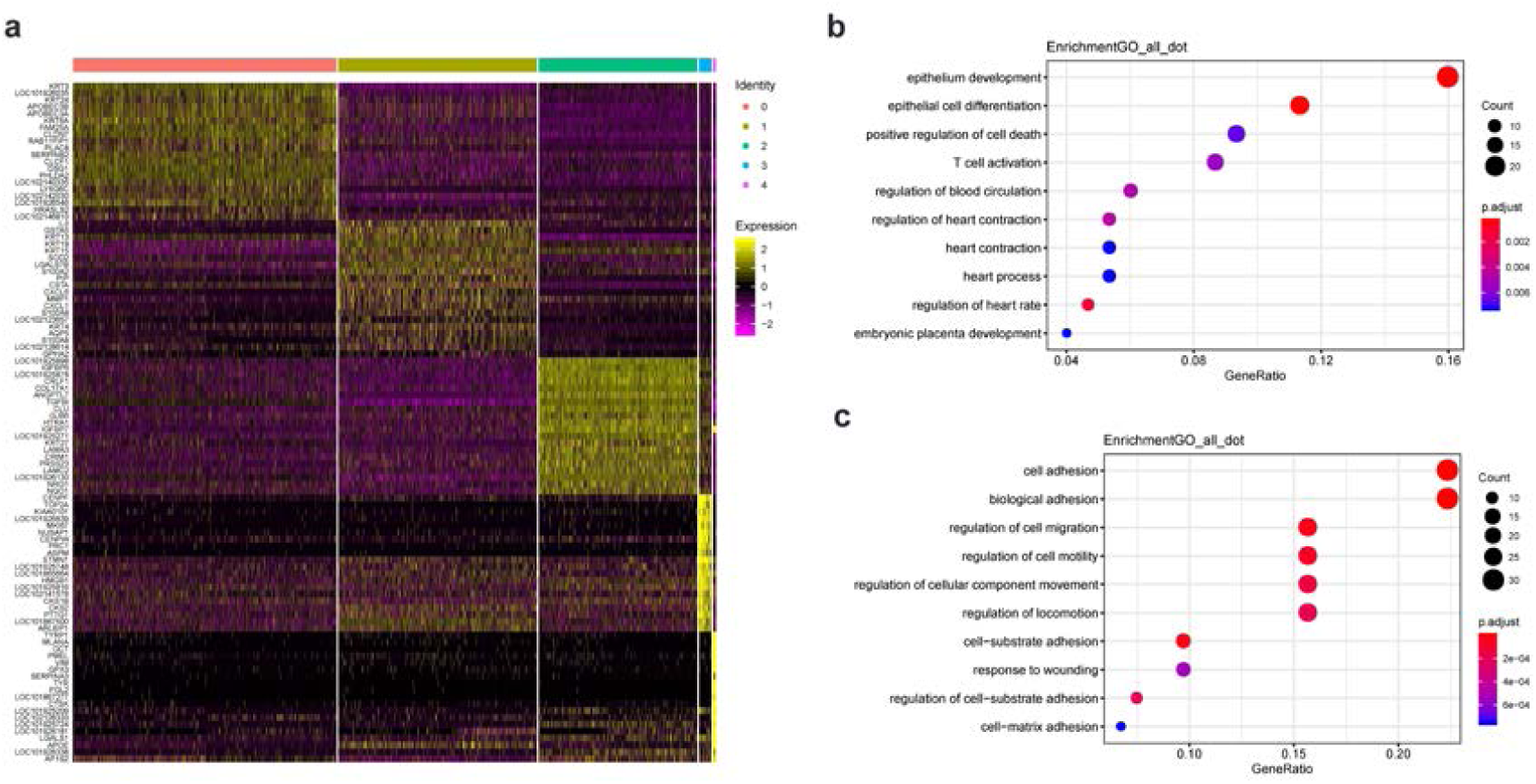
Cell-type and marker-gene identification in the crab-eating monkey cornea. **a**, Heatmap shows the top 20 marker genes expressed in LEC, CEC and TAC respectively. Color bars on the top are used to discriminate different cell types. **b**, GO items enriched in the genes up-regulated in CEC compared to TAC. **c**, GO items enriched in the genes down-regulated in CEC compared to TAC.

**Supplementary figure 8.**
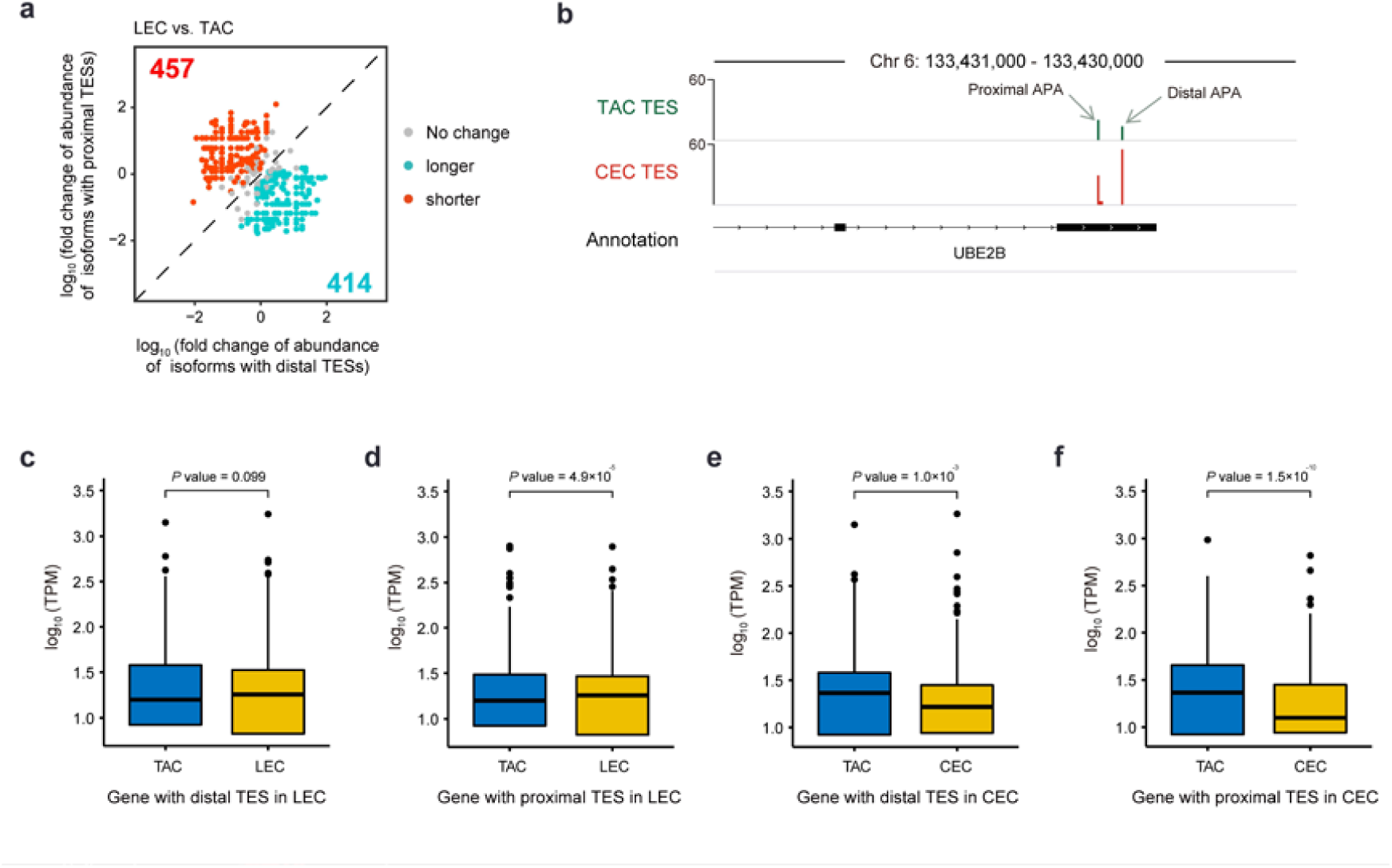
Differences between LEC and TAC in terms of RNA expression and APA choices. **a**, Expression data with isoform specificity reveals isoform expression differences between LEC and TAC (n = 956 genes). **b**, Genome browser track shows an example of APA choices for the gene *UBE2B* during differentiation of CEC from TAC (n = 414 genes). **c**, Boxplot comparing expression of genes in LEC and TAC, which have distal TESs in LEC (n = 414 genes). **d**, Boxplot comparing expression of genes in LEC and TAC, which have proximal TESs in LEC (n = 457 genes). **e**, Boxplot comparing expression of genes in CEC and TAC, which have distal TESs in CEC (n = 244 genes). **f**, Boxplot comparing expression of genes in CEC and TAC, which have proximal TESs in CEC (n = 285 genes). For **c-f**, significance was computed using two-sided Wilcoxon test.

**Supplementary table 1.**

| Sample | Sequencing depth |
| --- | --- |
| ERCC_01 | 1,052,270 |
| ERCC_02 | 1,571,949 |
| ERCC_03 | 3,799,426 |
| ERCC_04 | 3,989,246 |
| ERCC_05 | 2,291,544 |
| ERCC_06 | 3,835,792 |
| ERCC_07 | 3,986,186 |
| ERCC_08 | 7,964,775 |
| ERCC_09 | 7,645,733 |
| ERCC_10 | 5,223,629 |
| Average | 4,136,055 |

Sequencing depth for each ERCC spike-in libraries generated by scCAT-seq methods are listed.

**Supplementary table 2.**

| Library name | Sequencing Depth | TSS number | TES number | Gene number |
| --- | --- | --- | --- | --- |
| D41_71 | 2,493,578 | 11,155 | 4,867 | 3,917 |
| D44_52 | 3,100,138 | 12,906 | 5,667 | 4,594 |
| D44_72 | 1,612,099 | 10,693 | 4,601 | 3,558 |
| D45_52 | 2,723,321 | 12,858 | 4,866 | 4,066 |
| D45_72 | 2,080,059 | 11,355 | 4,352 | 3,570 |
| D46_71 | 3,955,396 | 13,570 | 5,547 | 4,674 |
| D47_52 | 1,628,466 | 10,786 | 5,433 | 4,295 |
| D47_72 | 1,056,585 | 9,567 | 4,276 | 3,232 |
| D48_52 | 1,484,463 | 10,165 | 4,180 | 3,346 |
| D48_72 | 1,869,191 | 10,710 | 4,407 | 3,501 |
| D49_72 | 1,330,893 | 9,587 | 3,671 | 2,902 |
| D50_52 | 1,518,348 | 10,201 | 4,288 | 3,474 |
| D50_71 | 2,249,770 | 10,802 | 4,104 | 3,343 |
| D50_72 | 2,717,055 | 11,725 | 5,136 | 4,262 |
| D51_52 | 3,733,854 | 12,258 | 5,437 | 4,600 |
| D51_72 | 2,474,653 | 11,314 | 3,860 | 3,253 |
| D52_52 | 2,624,766 | 12,142 | 5,692 | 4,689 |
| D52_72 | 4,015,130 | 13,347 | 5,869 | 4,998 |
| Average | 2,370,431 | 11,397 | 4,792 | 3,904 |

Information about 18 single DRG neurons generated by scCAT-seq method for further benchmarking in this study are listed in this table. A gene is detected only if there are TSS peaks and TES peaks mapped to the said gene.

**Supplementary table 3.** ScISOr-Seq information in this study.

| Sample | Subreads<br>base (G) | Number of<br>Subreads | Average subreads<br>length | Number<br>of CCS | Number<br>of FLNC |
| --- | --- | --- | --- | --- | --- |
| DRG_1 | 0.32 | 214,295 | 1,487 | 19,947 | 4,289 |
| DRG_2 | 0.35 | 237,777 | 1,481 | 23,070 | 3,835 |
| OC_1 | 0.89 | 612,741 | 1,446 | 47,152 | 8,171 |
| OC_2 | 0.24 | 165,171 | 1,436 | 13,305 | 780 |
| OC_3 | 1.03 | 703,398 | 1,462 | 54,258 | 22,549 |
| OC_4 | 3.52 | 2492,563 | 1,414 | 193,637 | 13,932 |
| OC_5 | 0.22 | 145,585 | 1,483 | 14,857 | 1,009 |
| OC_6 | 1.31 | 912,567 | 1,441 | 68,471 | 27,559 |

Information about 6 single oocytes and 2 single DRG neurons generated by scISO-seq are listed.

**Supplementary table 4.**
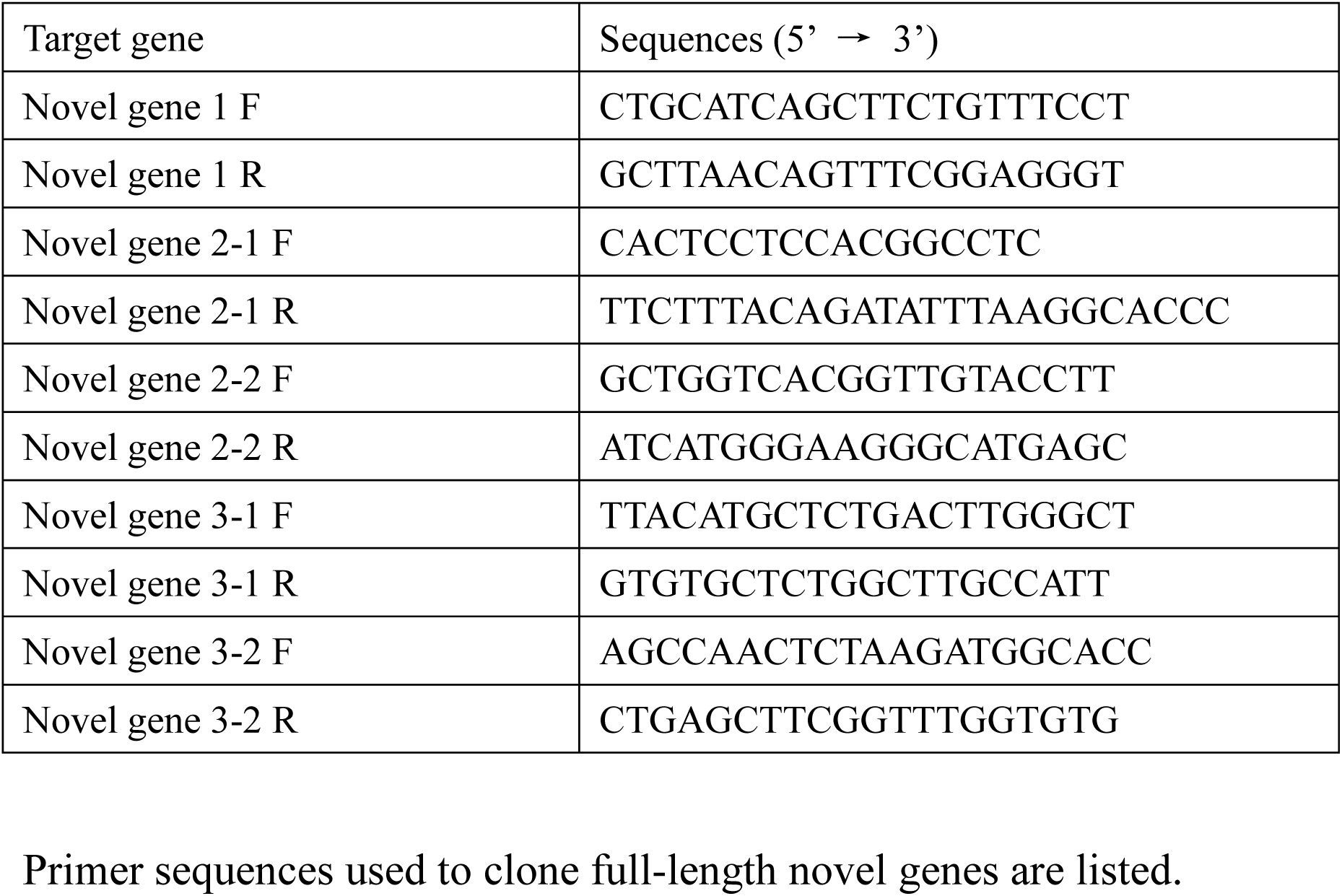
Cloning primers used in this study.

## References

1. Trapnell C (2015) Defining cell types and states with single-cell genomics. Genome Res 25(10):1491–1498.

2. Wagner A, Regev A, & Yosef N (2016) Revealing the vectors of cellular identity with single-cell genomics. Nat Biotechnol 34(11):1145–1160.

3. Tang FC, et al. (2009) mRNA-Seq whole-transcriptome analysis of a single cell. Nat Methods 6(5):377–U386.

4. Regev A, et al. (2017) The Human Cell Atlas. Elife 6.

5. Noseda M & Harding SE (2018) Understanding dynamic tissue organization by studying the human body one cell at a time: the human cell atlas (HCA) project. Cardiovasc Res 114(12):E93–E95.

6. Barash Y, et al. (2010) Deciphering the splicing code. Nature 465(7294):53–59.

7. Pan Q, Shai O, Lee LJ, Frey J, & Blencowe BJ (2008) Deep surveying of alternative splicing complexity in the human transcriptome by high-throughput sequencing. Nat Genet 40(12):1413–1415.

8. Donczew R & Hahn S (2018) Mechanistic Differences in Transcription Initiation at TATA-Less and TATA-Containing Promoters. Mol Cell Biol 38(1).

9. Di Giammartino DC, Nishida K, & Manley JL (2011) Mechanisms and consequences of alternative polyadenylation. Mol Cell 43(6):853–866.

10. Moqtaderi Z, Geisberg JV, & Struhl K (2018) Extensive Structural Differences of Closely Related 3’ mRNA Isoforms: Links to Pab1 Binding and mRNA Stability. Mol Cell 72(5):849–861 e846.

11. Reyes A & Huber W (2018) Alternative start and termination sites of transcription drive most transcript isoform differences across human tissues. Nucleic Acids Res 46(2):582–592.

12. Liu YL & Elliott DJ (2010) Coupling genetics and post-genomic approaches to decipher the cellular splicing code at a systems-wide level. Biochem Soc T 38:237–241.

13. Anvar SY, et al. (2018) Full-length mRNA sequencing uncovers a widespread coupling between transcription initiation and mRNA processing. Genome Biol 19.

14. Chen W, et al. (2017) Alternative Polyadenylation: Methods, Findings, and Impacts. Genom Proteom Bioinf 15(5):287–300.

15. Gupta I, et al. (2018) Single-cell isoform RNA sequencing characterizes isoforms in thousands of cerebellar cells. Nat Biotechnol 36(12):1197-+.

16. Hochgerner H, et al. (2017) STRT-seq-2i: dual-index 5’ single cell and nucleus RNA-seq on an addressable microwell array. Sci Rep-Uk 7.

17. Kouno T, et al. (2019) C1 CAGE detects transcription start sites and enhancer activity at single-cell resolution. Nat Commun 10.

18. Goetz JJ & Trimarchi JM (2012) Transcriptome sequencing of single cells with Smart-Seq. Nat Biotechnol 30(8):763–765.

19. Picelli S, et al. (2014) Full-length RNA-seq from single cells using Smart-seq2. Nat Protoc 9(1):171–181.

20. Byrne A, et al. (2017) Nanopore long-read RNAseq reveals widespread transcriptional variation among the surface receptors of individual B cells. Nat Commun 8.

21. Ng P, et al. (2005) Gene identification signature (GIS) analysis for transcriptome characterization and genome annotation. Nat Methods 2(2):105–111.

22. Haberle V, Forrest ARR, Hayashizaki Y, Carninci P, & Lenhard B (2015) CAGEr: precise TSS data retrieval and high-resolution promoterome mining for integrative analyses. Nucleic Acids Res 43(8).

23. Islam S, et al. (2014) Quantitative single-cell RNA-seq with unique molecular identifiers. Nat Methods 11(2):163-+.

24. Arguel MJ, et al. (2017) A cost effective 5’ selective single cell transcriptome profiling approach with improved UMI design. Nucleic Acids Res 45(7).

25. Consortium F, et al. (2014) A promoter-level mammalian expression atlas. Nature 507(7493):462–470.

26. Wang R, Nambiar R, Zheng D, & Tian B (2018) PolyA_DB 3 catalogs cleavage and polyadenylation sites identified by deep sequencing in multiple genomes. Nucleic Acids Res 46(D1):D315–D319.

27. Zeisel A, et al. (2015) Cell types in the mouse cortex and hippocampus revealed by single-cell RNA-seq. Science 347(6226):1138–1142.

28. Hashimshony T, et al. (2016) CEL-Seq2: sensitive highly-multiplexed single-cell RNA-Seq. Genome Biol 17:77.

29. Velten L, et al. (2015) Single-cell polyadenylation site mapping reveals 3’ isoform choice variability. Mol Syst Biol 11(6).

30. Leung MK, Xiong HY, Lee LJ, & Frey BJ (2014) Deep learning of the tissue-regulated splicing code. Bioinformatics 30(12):i121–129.

31. Qin Z, Stoilov P, Zhang X, & Xing Y (2018) SEASTAR: systematic evaluation of alternative transcription start sites in RNA. Nucleic Acids Res 46(8):e45.

32. Hu YJ, et al. (2016) Simultaneous profiling of transcriptome and DNA methylome from a single cell. Genome Biol 17.

33. Liao Y, Smyth GK, & Shi W (2014) featureCounts: an efficient general purpose program for assigning sequence reads to genomic features. Bioinformatics 30(7):923–930.

34. Kharchenko PV, Silberstein L, & Scadden DT (2014) Bayesian approach to single-cell differential expression analysis. Nat Methods 11(7):740–U184.

35. Li H (2018) Minimap2: pairwise alignment for nucleotide sequences. Bioinformatics 34(18):3094–3100.

36. L. B (2001) Random forests. Machine learning 45(1):5–32.

37. Breiman L, Friedman, J.H., Olshen, R.A., and Stone, C.I. (1984) Classification and regression trees. (Belmont, Calif.: Wadsworth).

38. Boser BE, Isabelle M. Guyon, and Vladimir N. Vapnik. (1992) A training algorithm for optimal margin classifiers. Proceedings of the fifth annual workshop on Computational learning theory. ACM, pp 144–152.

39. Dietterich TG (2000) Ensemble methods in machine learning. in International workshop on multiple classifier systems (Springer, Berlin, Heidelberg).

40. Quinlan AR & Hall IM (2010) BEDTools: a flexible suite of utilities for comparing genomic features. Bioinformatics 26(6):841–842.

41. Stuart T, et al. (2019) Comprehensive Integration of Single-Cell Data. Cell 177(7):1888–1902 e1821.

42. Trapnell C, et al. (2014) The dynamics and regulators of cell fate decisions are revealed by pseudotemporal ordering of single cells. Nat Biotechnol 32(4):381–386.

